# Computational Modeling: Human Dynamic Model

**DOI:** 10.1101/2020.08.23.262048

**Authors:** Lijia Liu, Joseph L. Cooper, Dana H. Ballard

## Abstract

Improvements in quantitative measurements of human physical activity are proving extraordinarily useful for studying the underlying musculoskeletal system. Dynamic models of human movement support clinical efforts to analyze, rehabilitate injuries. They are also used in biomechanics to understand and diagnose motor pathologies, find new motor strategies that decrease the risk of injury, and predict potential problems from a particular procedure. In addition, they provide useful constraints for underlying neural circuits. This paper describes a physics-based movement analysis method for analyzing and simulating bipedal humanoid movements. A 48 degree of freedom dynamic model of humans has been developed to report humanoid movements’ energetic components. It has sufficient speed and accuracy to analyze and synthesize real-time interactive applications, such as psychophysics experiments using virtual reality or human-in-the-loop teleoperation of a simulated robotic system. The dynamic model is fast and robust while still providing results sufficiently accurate to be used to believably animate a humanoid character or estimate internal joint forces used during a movement for creating effort-contingent experimental stimuli. A virtual reality environment developed as part of this research supports controlled experiments for systematically recording human behaviors.

## Introduction

The complexity of human motion was first dramatically captured via the Muybridge high-speed photographs [1] which spawned a number of separate analysis techniques in different disciplines. Visualization first used keyframing techniques but later sophisticated models used in advanced rendering for computer graphics e.g. [2]. The early cognitive analyses of human behavior [3] focused on human motion in problem-solving, using an essentially logical approach. In robotics, sights have been obtained by building physical systems directly [4] that straddle the boundary between humans and robotics that have shed light on the human design. However, these efforts are characteristically specialized. In another development, machine learning techniques have been introduced for use in analyzing animal-like motion [5].

Most recent advances in the speed of computing and novel formulations of the dynamic equations of motion have engendered a new approach to understanding human movement fundamentals. Large scale human movement models can be built with the objective of understanding how the human generates goal-oriented behaviors in real-time. However, modeling all the complexity of the human musculoskeletal system can be daunting, with over 600 muscles controlling a complex skeletal system with over 300 degrees of freedom. Moreover, to control this complexity, in addition to its vast cortical memory system, the forebrain coordinates specialized subsystems such as the Basal Ganglia and Thalamus in realizing human real-time movement coordination. The upshot is that progress tends to be specialized [6], and there are many open problems [7].

In the face of these complex challenges, a major alternate modeling route is to forego the neural level of detail as well as one that features muscles and model more abstract versions of the human system that still use multiple degrees of freedom but summarize muscle effects through joint torques. The computation of the dynamics of such multi-jointed systems recently has also experienced significant advances. The foremost of these, use a kinematic plan to integrate the dynamic equations directly. Several different systems exist, such as MuJoCo, Bullet, Havok, Open Dynamic Engine(ODE)^1^, and PhysX, but an evaluation by [8] found them roughly comparable in capability, and only MuJoCo^2^ has been applied to human modeling.

Thus there is a need for an exclusively human movement based model that could be used to inform laboratory experiments [9], clinical studies e.g [10] also verify experiments that have only qualitative results [11, 12]. Our human dynamic model (HDM)^3^ has a singular focus on human movement modeling and uses a unique approach to integrating the dynamic equations. A direct dynamics integration method to extracts torques from human subjects in real-time [13–15] using a unifying spring constraint formalism.

The HDM system is built on top of the physics engine ODE, but has two significant innovations added in order to handle the closed-loop kinematic chains of bipedal movements and the contact constraints they introduce, which have proven difficult to model. One is to allow the kinematic makers of a motion capture system to be modeled as very large point masses. The result is to stabilize the integration of the underlying dynamic equations. The other is to allow the reduction of contact constraints into stiff springs, which has the result of allowing the incorporation of external forces and points of contact.

Most of the computation of the joint torques uses the kinematic data, but there is a balance issue to be dealt with.For example human motions for familiar tasks such as balancing while putting on socks can use on remembered protocols, if they can depend ancillary system such as the vestibular to correct errors.In the same way we use use a similar closed loop system to generate small corrections.

The focus of the paper is to describe the HDM simulator as a useful laboratory instrument as well as describe demonstrations that lend support to the kinematic plan approach to movement memory. These goals are illustrated and evaluated in several different demonstrations to illustrate the versatility of the method.

## Model Overview

The HDM is a fast, robust, intuitive, and inexpensive multi-purpose tool for simulating, analyzing, and synthesizing humanoid movement. Fig. 1 shows a frame from a study of the cost of movements used in a virtual tracing experiment [16]. The model interface^4^ shown in allows the construction of the human model using the physics engine via a multi-purpose graphical interface for analyzing movement data captured through interaction with the virtual environment. With this tool, it is possible to interactively fit a model to marker data, dynamically adjust parameters to test different effects, and visualize the results of kinematic and dynamic analysis. Another example is shown in Fig 2, which shows frames from a jumping sequence made originally by a human subject and then recreated by the HDM system using the inverse dynamic method.

**Fig 1.**
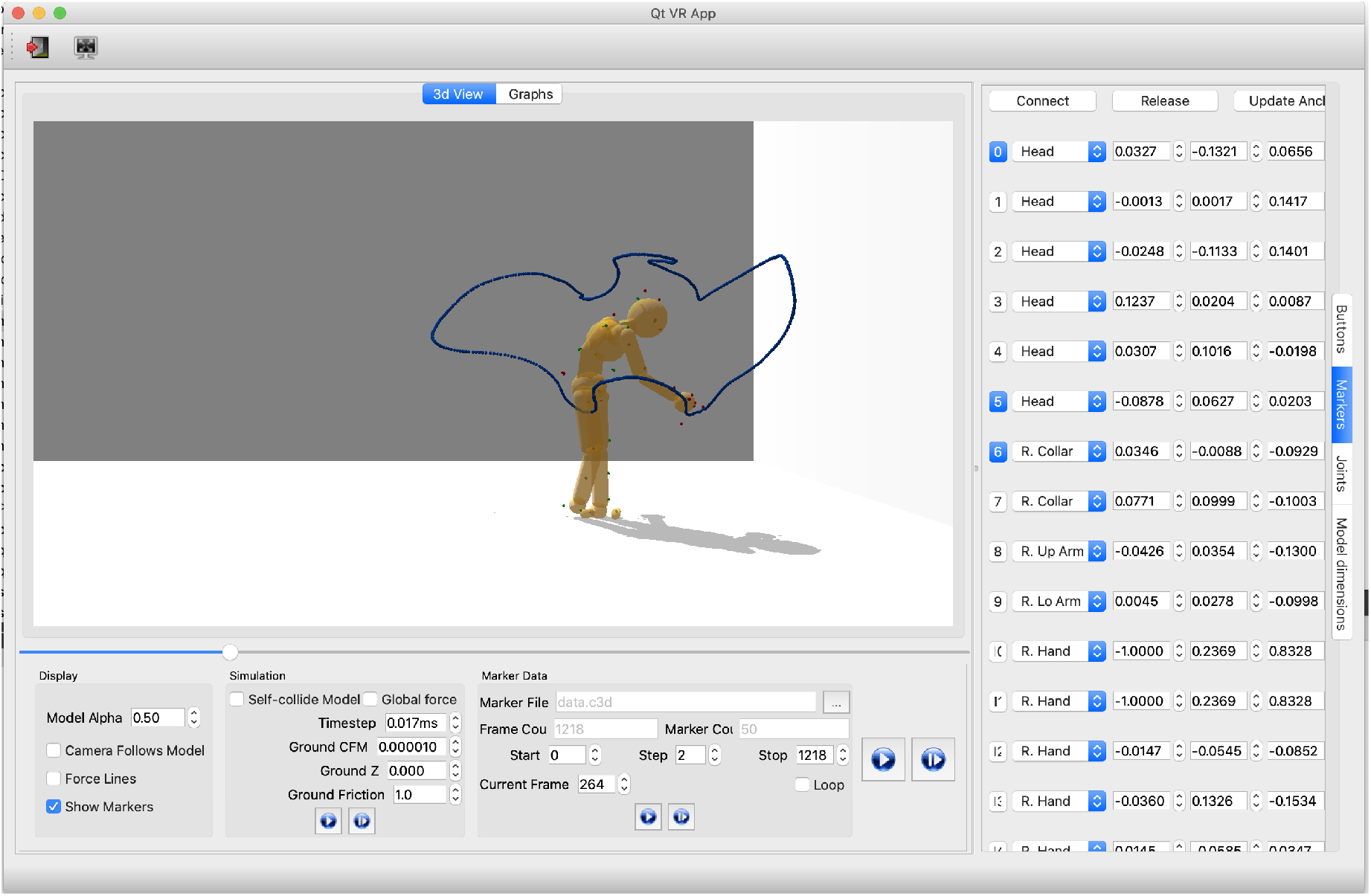
The HDM user interface. It supports various visualizations of relevant data and control for analyzing and producing physically-based movements. The programmed parameters of the model consist of physical world parameters, joints constraints, and the model’s body-marker relative positions. In this depiction shows how users can get the current HMD configurations by clicking the buttons on the rightmost vertical menu.“Marker” is selected, meaning the marker information is shown:(1) The first column represents marker index buttons. Buttons in blue means the corresponding markers are attached to the HDM. Users can attach/detach markers by clicking index buttons. (2) The second column shows body segments where markers are attached. Each spin box is a collective item of all body segment names. Users can use it to change the body-marker attachment relationship. (3) The three-five columns present the marker-body relative positions. Users can modify the values directly using this interface. (4) The “Connect” button and “Release” button on the top are to attach or detach all the markers, respectively. The “Update Anchor” button automatically updates the marker-body relative positions based on the current motion posture.

**Fig 2.**
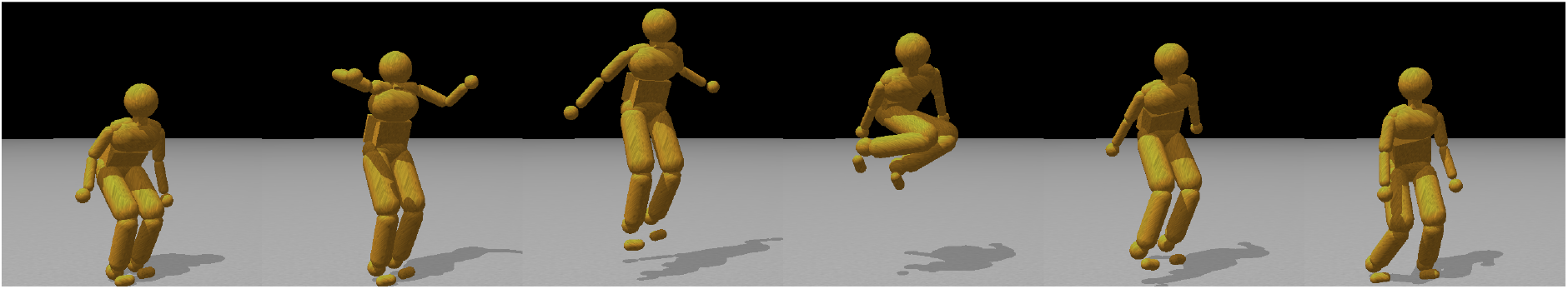
Model capability illustration. A jump sequence reproduced with physics-engine-based inverse dynamics using recorded motion capture data from a human subject. The recreated jump height is achieved completely from ground forces, with small residual torques (≤ 100*Nm*) keeping the model from tipping over.

Analysis of human motions utilizing the human dynamic model is implemented in the following five steps:

1. Motion synthesis: it simulates human motion by following the motion capture data [14]
2. Inverse kinematics: it calls the ODE built-in functions to compute the joint angles and joint angular velocities at each frame.
3. Forward kinematics: it simulates human motion based on the computed joint properties. This step is to check the correctness of recovered kinematic properties.
4. Inverse dynamics: it calls the ODE built-in function to compute the required joint torques.
5. Forward dynamics: it simulates human motion based on the computed torques/forces. This step is to check the correctness of recovered dynamic properties.

At each frame, instantaneous power was computed from the product of net joint torque and joint angular velocity. The work performed at each joint were determined by numerically integrating the instantaneous powers over the entire tracing task. In this way, the the energy cost of human motions can be computed given motion capture data.

More details of building the HDM are described in the Method section. The derivation of the mathematics underlying the physics simulation is presented separately in S1 Appendix.

This section focuses on describing the model’s capabilities through a series of examples in different settings. Several test experiments provide qualitative and quantitative validation of the physics-based movement analysis techniques described here.

### Test 1: Model Performance

Given that the torque recovery technique will be the basis for our experiments, it is essential to establish its accuracy in absolute terms. A straightforward to do this is to use a *particular model* to generate joint torque data and then verify that these generating torques can be recovered with sufficient accuracy. To test the model accuracy and noise sensitivity, we first use the PhaseSpace motion capture system to gather the walking data and then let the model simulate the walking motion. To simulate possible sensor errors in the PhaseSpace system, we introduce noise into the simulated marker positions and study the accuracy of recovery with increasing noise levels.

#### Noise tolerance

Inverse dynamics computations rely on first finding the model’s pose. Therefore, given motion capture data, it is essential to synthesize the pose sequence precisely. We used the HDM to synthesize treadmill walking and then compute its accuracy. The aim of this study was to assess the effect of sensor noise on the results and compare the joint angles and torques found with our method to those used to generate marker data. We used an experimental process similar to that employed in [17]. In this experiment, both steps were tested by studying eight steps of marker data captured from treadmill walking. The movement lasts a little longer than 4 seconds, giving us 260 frames of data. For this computation, we used data sampled at 60 Hz.

We used a preliminary pass through the motion capture data to generate synthesized “ground truth” marker, pose, and torque data. After using the physics-based inverse kinematics to compute joint angles, we constrained the body to use forward dynamics to reproduce the joint angles with internal torques (and residual forces at the waist segment). As the model performed the movement, we recorded the global position of the marker attachment points. We also recorded the forces used and the resulting joint angles. Thus we had synthetic “ground truth” data directly from the model.

Using the synthetic marker data, we analyzed the process by perturbing all marker positions at each frame in time along all three axes with mean-centered Gaussian noise of a controlled standard deviation. Applying physics-based pose-fitting followed by inverse dynamics produced a new set of virtual marker positions, joint angles, and torques. The results are shown in Fig 3.

**Fig 3.**
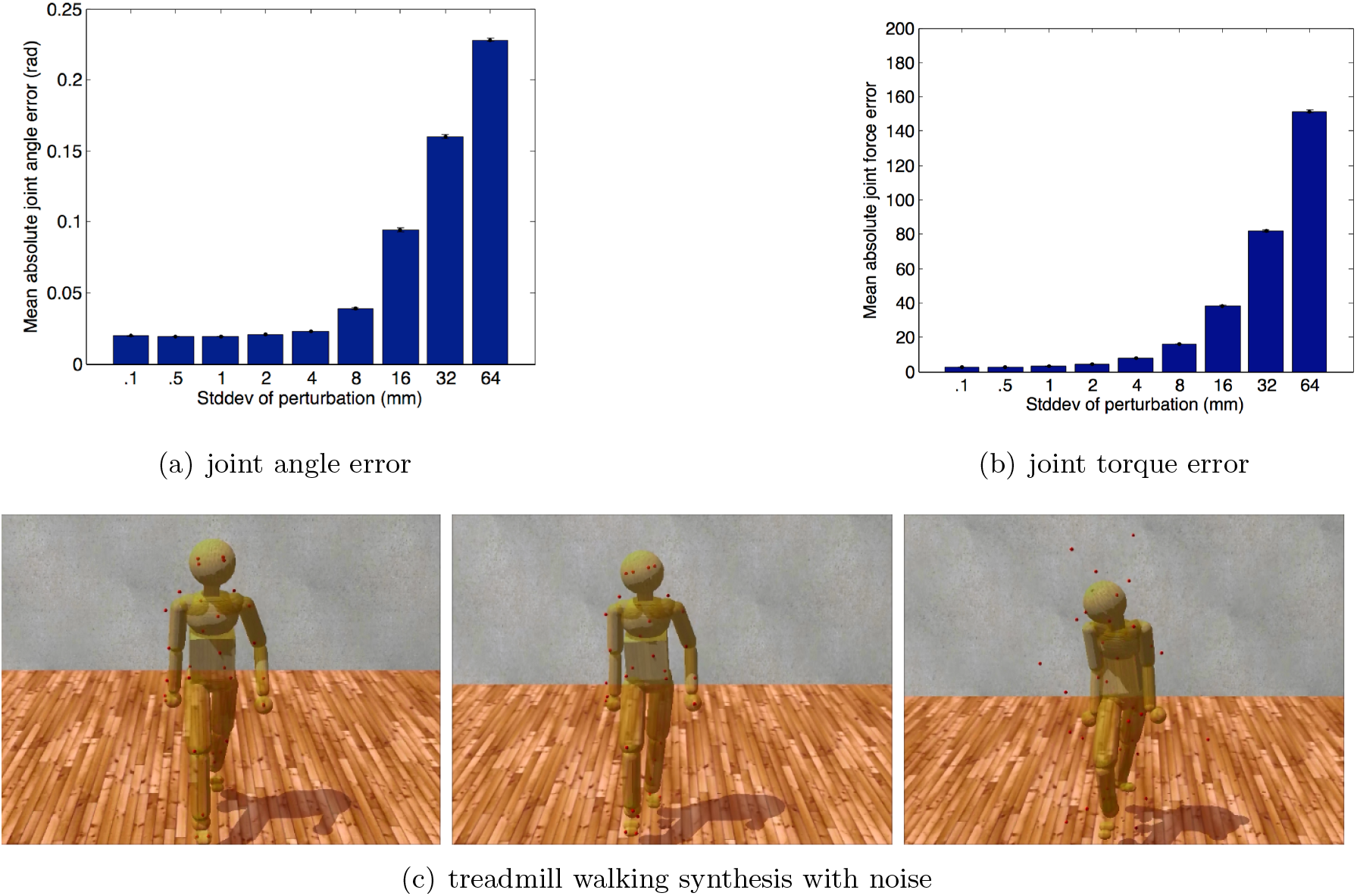
Model noise sensitivity. Error of joint angles, and internal torques resulting from physics-based inverse kinematics and inverse dynamics used to analyze perturbed marker data. We repeated the process twenty times for each noise-level at nine different standard-deviations. Standard-deviations, in mm, were (0.1, 0.5, 1, 2, 4, 8, 16, 32, 64). Error bars show standard error of the mean.(a) The accuracy of the PhaseSpace motion capture device is approximately 5mm over its 3 × 6 meter workspace, resulting an average angular error of 1 degree. (b) The same estimates for torque error are between 5 and 10 Nm, typically approximately 1%. These small errors are well within the requirements for our experiments. (c) Poses generated by forward dynamics using forces obtained from three inverse dynamics simulations based on Gaussian perturbed walking data (0.1mm, 8mm, and 64mm noise levels). Although at very high levels of noise, the model follows the reference motion poorly, the movement still looks, qualitatively, like walking.

Gaussian perturbations render the marker data dynamically inconsistent. This dynamic inconsistency also pushes a constrained system toward singularity, making it more challenging to solve numerically. We included very high levels of noise to see if they would slow the system down, or prevent it from finding any solution. In all cases, the system analyzed the perturbed data in real-time, finding pose data and dynamics data to fit the marker data.

After running through an inverse kinematics pass, an inverse dynamics pass, and a forward dynamics pass for each trial run; we compared the marker attachment points, joint angles, and joint torques from the forward dynamics pass to the synthetic ground truth data. Fig. 3 shows the mean error for across all degrees of freedom and frames of time for each quantity measured. Although the perturbations make the marker data dynamically inconsistent, small amounts of noise have minimal effect on the computed measurements. Fig 3 shows that functional recovery is possible with up to 8mm standard error deviations. A ±1mm PhaseSpace marker position accuracy translates in our model into an average joint angle error of 0.02 radians and average force errors of 3 Newtons.

There is a systematic error in both the marker positions and joint angles caused by the fact that the constraints behave like springs. The spring-like behavior causes the marker positions and joint angles to lag behind their targets by a small amount and dampens the overall movement. This lag and damping are apparent in Fig. 4 comparing individual trajectories for selected dimensions of the joint angles and torques. As shown in Fig 4, the data follow ground truth very well under low noise conditions.

**Fig 4.**
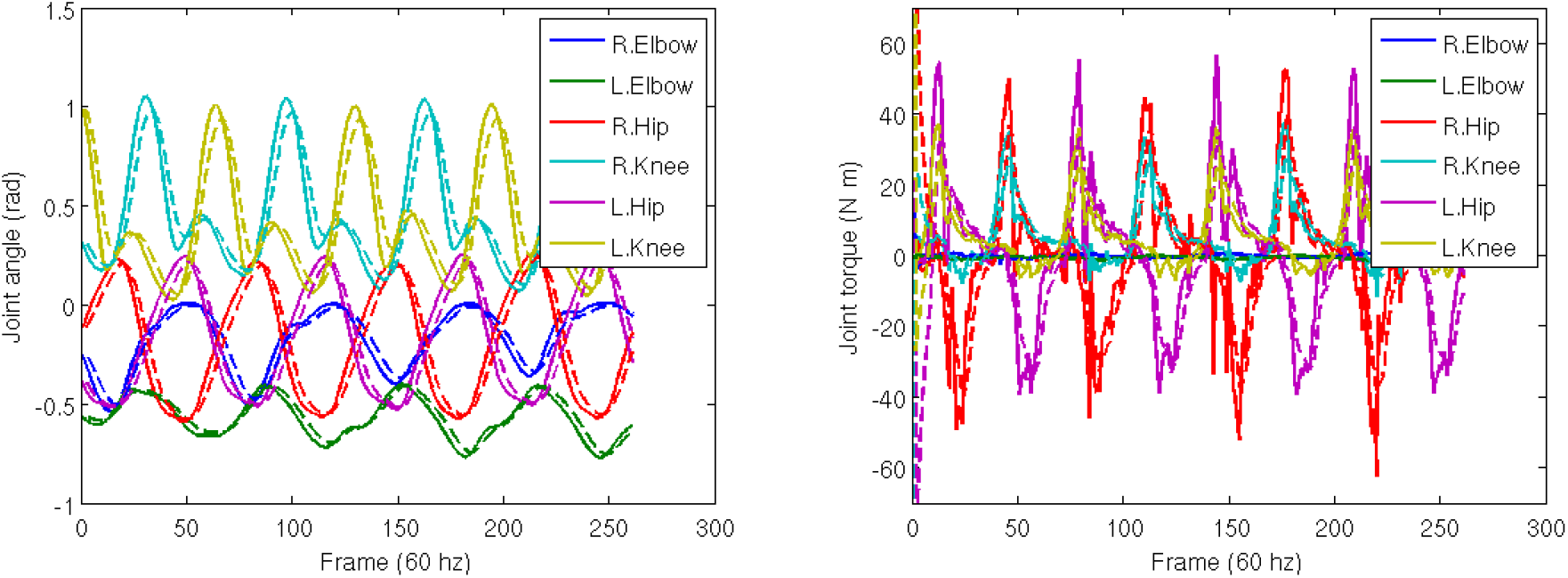
Trajectory reconstruction. Trajectories of selected degrees of freedom from the perturbation study. Solid lines show ground truth. Dashed lines show computed data. Simulated spring forces make the computed data lag behind and smooth the ground truth.

#### Residual torques/forces and ground forces

The inverse dynamics uses measured kinematics and external forces to calculate net joint torques in a rigid body linked segment model. [18] However, discrepancies between the dynamic forces of the model and the kinematic of the reality make it so that the dynamic model falls over unless action is taken to stabilize it. Adjustments to internal joint torques can be used to stabilize the body but cause the body’s pose to deviate from its intended pose. A common way to compensate this problem is by introducing “residual forces and torques.” In humans, these additions would be consequential of measurements inthe human vestibular system. The HDM includes a joint to the model’s waist to constrain it to reproduce orientation deviations found during the pose-fitting pass. To minimize the effect of these external forces, we used torque limits on the amount of stabilizing torque available.

The system fully configured system could be tested against an objective set of measurements. We compaired HDM data together with ground force data from a pair of balance boards. Fig 5 shows the calibration of the ground force computed from our method compared to those taken from *Wii^T^ ^M^* force plates. A subject standing on two force plates, varied their stance from one being supported exclusively by leg standing on one plate and then shifted their weight to the other leg to be supported by the other plate. For this simple movement of transitioning from standing on one foot or the other, residual angular torques of 30Nm were sufficient to keep the dynamic model quite close to its target trajectory.

**Fig 5.**
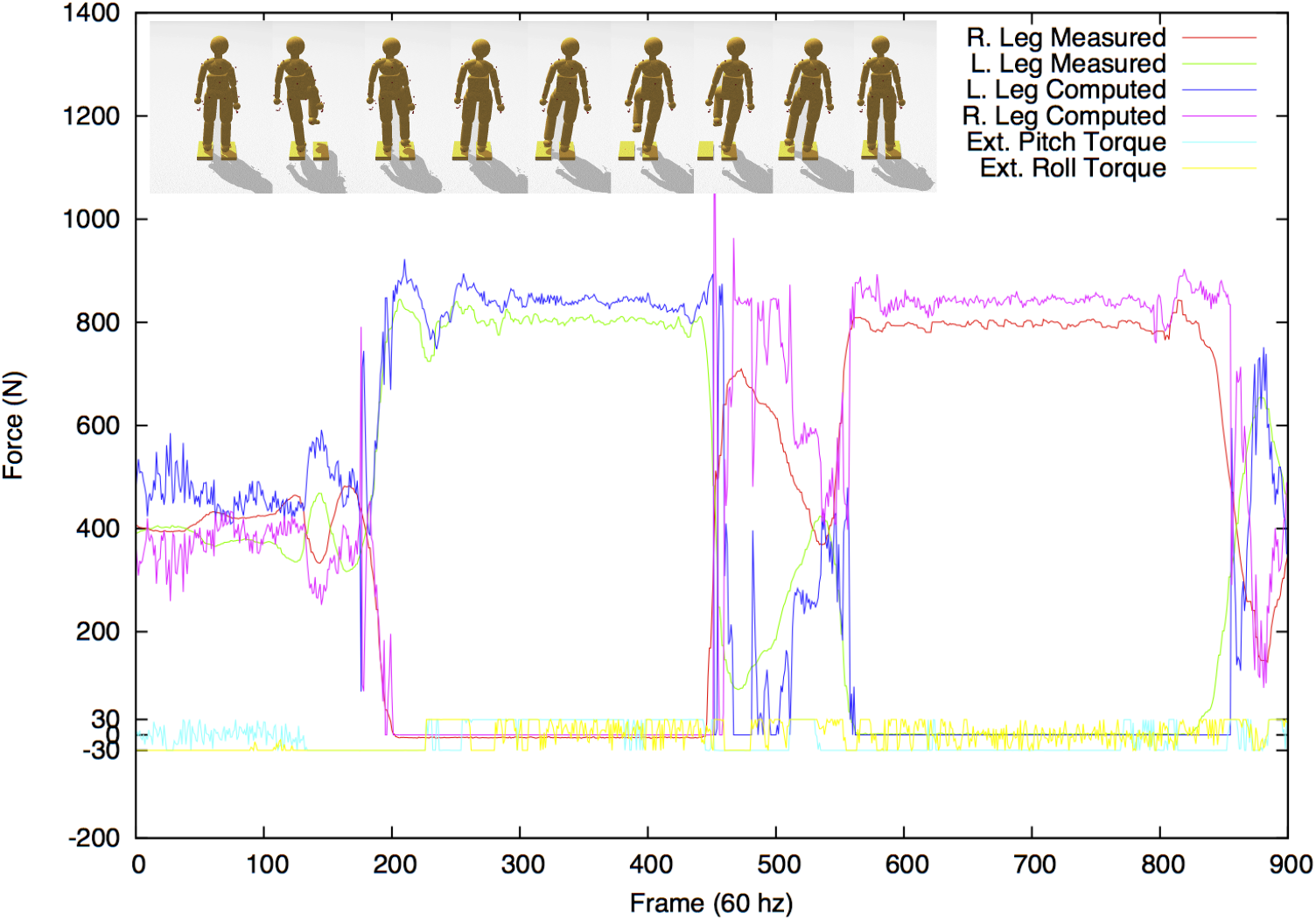
Comparing ground forces between the model and the Wii force plate. (Top) Two Wii force plates serve as accurate calibration reference. A subject stood on the two plates and then changed stances, balancing first on the left foot and next on the right. (Bottom) The comparison between the measurement systems is surprisingly good, during the stance phases, showing only a 10% difference between the *measured* ground forces and the *computed* forces.

The residual torques are very modest, being within ±5% of the maximum excursion. The correspondence is actually a little better as the faux vestibular balance forces are not factored into the comparison. Note also that we cannot expect the correspondence to be exact during the phase between the two stances as there is no attempt in the model in this test to make the dynamics of the changing stance match that of the force plates. To generate independent movements, such as grasping might need additional accuracy [19], but for estimating a subject’s energetic cost, the accuracy is well within range.

Fig 5 also shows the comparison results between the sensor-measured ground forces for the right and left feet (red and green lines) with the computed ground forces found through physics-based inverse dynamics (blue and pink lines). During bipedal stance phase, the forces come surprisingly close. The largest discrepancies come during the transition from one foot to the other. These discrepancies can be blamed largely on poor collision detection resulting from an abstract model of the foot.

### Test2: Model Validation

The previous demonstrations report on tests of the accuracy of the system in completely artificial situations. Herein we describe three tests of the whole body model’s ability to fit data obtained from human subjects. The first test uses a subject carrying out successively more difficult reaches in a virtual reality environment to test whether the model’s estimate of movement costs correlate with increasing task difficulty. The second test simulates data from an issue facing movements in an aging population. Do aging subjects’ reduced use of arm swing while walking incur a movement cost, and does the HDM’s estimate correspond to laboratory treadmill data? The final test demonstrates an essential property of the model concerning its degrees of freedom. The critical observation is that virtues of their interconnections constrain the degree of freedom of the model; thus, the control of a posture can be achieved with a very reduced set of key marker positions. This has implications for movement control programs.

#### Whole body reaching

The movement accuracy test is encouraging, but the importance of the method depends on its usefulness to capture the energetic cost of whole-body movements in a complex experimental setting. One such venue is a three-dimensional Virtual Reality (VR) environment. The advantage of the VR environment for studying human movements is that the dimensions and the dynamic variations of the parametric quantities describing the setting can be varied with full experimental control.

In this experiment, we studied where human subjects needed to use whole-body movements cost choosing actions. From a particular start, a human subject touched targets suspended in 3D space. The experimental setup is demonstrated in Fig. 6. The subject is wearing the PhaseSpace motion capture suit and the nVisor head-mounted stereo display. From a fixed starting position, a subject is instructed to touch one of the targets and return to the starting position.

**Fig 6.**
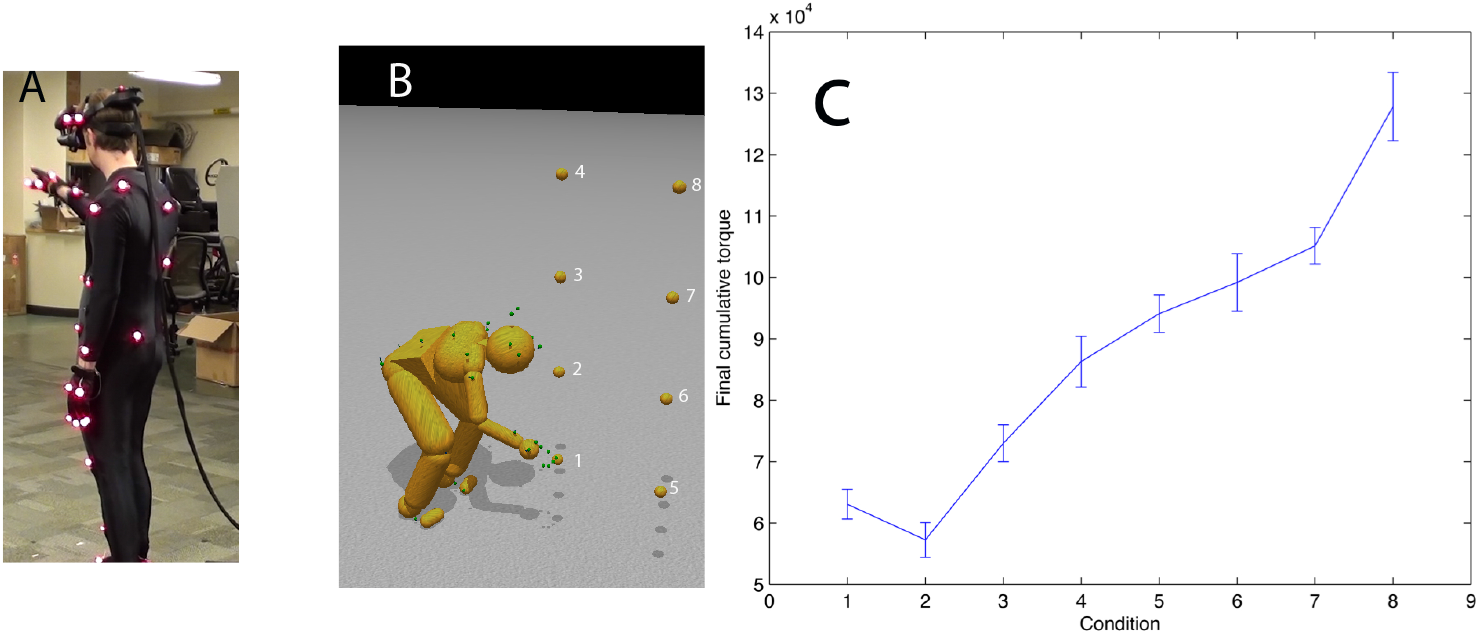
Reaching in a virtual reality environment. A) A subject reaches to touch virtual targets seen in a HMD. The Ss’ reach is unconstrained. B) The subject reaches to the different numbered targets on separate trials. C) The average integrated torque over 10 trials per reach shows that the method reliably discriminates between movement costs for the further and higher locations.

Tests were able to establish that, just focusing on integrated net torque and avoiding stiffness, the total cost of a movement recorded by our system reliably discriminates the energetic costs of the movement in the way hypothesized. The hypothesized cost of reaching for and touching each of the targets was ranked on the basis of distance and height relative to the subject. Note that target 2 is the least expensive as the subject does not have to crouch or extend significantly to touch it. Targets 5 through 8 are more costly than targets 1 through 4 as they require that the subject take a step to touch them. These results were expected, but the point was to show that the overall setting and model could produce reliable torque estimates.

This demonstration shows that the model can be used in any setting where the cost of a movement is hypothesized to be a constituent factor. We develop this technique further in the next demonstration.

#### Comparing the HDM with a prior experimental result

Once the joint stiffness parameters were adjusted appropriately, can it reproduce the results of a stiffness modulating experiment? The experiment we tried was to replicate that of Ortega et al. [20]. They showed that arresting the arm swing during treadmill walking incurred an increased metabolic cost of 6%. Our hypothesis was that to reproduce this result we could modify our walking data for the model so that the arms were clamped by the sides with stiff stationary markers.

To test this feasibility, we used one of our HDM walking data sets in a test situation. The cost of walking was computed and with a modification designed to model the data in [20]. To simulate their experiment, we modified the model data so the arms could swing with the walking gait for the standard case, but for the restricted case, the arms were constrained by markers that move with the stride but are not allowed to swing. Since the arms under restricted situation were not allowed to balance the leg movements, we expected the energetic cost to be higher. As shown in Figure 7, the result was that the constrained walk was about 6 % more expensive than the standard walk, which was essentially the value obtained by the Farley lab [20]. The use of the HDM in imitating this experiment shows off the utility of the model; no elaborate tuning was necessary to obtain the preliminary result other than restraining the arms.

**Fig 7.**
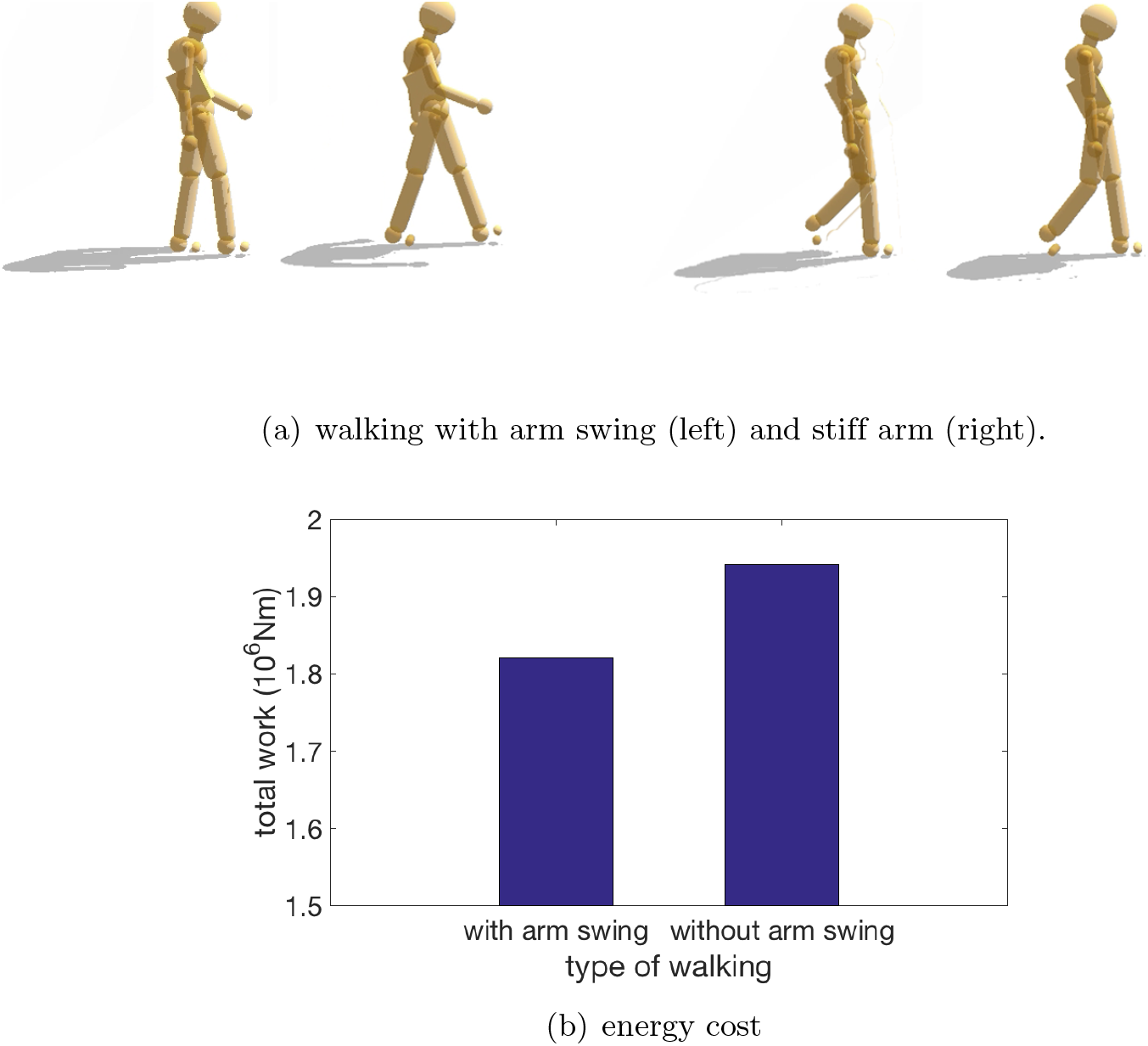
Comparison of efforts while walking with/without arm swing. (a) In a preliminary test of our design, the energetic cost of normal walking is compared to the case where the arms are constrained from swinging. Our hypothesis is that if subjects are instructed to walk without moving their arms, they will accomplish this by using muscle co-contraction and that this effect can be realized in the HDM with stationary markers that keep the arms vertical. (b)The increased cost measured by the HDM is 6.1 %, extremely close to the 6 % result obtained by Ortega [20].

#### Controlling poses using reduced marker sets

Tests of movement accuracy revealed that the dynamics engine was able to tolerate significant noise levels added to the marker positions. Another possibility is to use a subset of the markers to constrain the dynamics and still produce reasonable walking gaits. Human pose sequences from simple single-behavior motions lie on a very low-dimensional linear subspace [21]. However, original feature space of human motions has two many dimensions, e.g. the HDM uses 51 markers, so one pose is represented by a 123-dimension coordinate system. Tests show that for many movements, with suitable internal stiffness, it is only necessary to control the location of a reduced set consisting of the head, hands, and feet markers [22]. This property could have been expected from studies of muscle synergies, which show that muscle contractions coordinate in movement generation [23, 24].

Fig. 8 shows a qualitative comparison between a pose found using the whole marker set (on the left) and one found using only head, hands, and feet(on the right). To achieve the reduced marker pose, we started the model in an upright stance with the arms by the side, and then the reduced set markers are moved slowly along trajectories that leave them in the final posture. The straight arms take advantage of the elbow joint angle limitation. Joint limits on the knees and elbows and general joint stiffness naturally bias the physics engine to find a pose that is very close to the fully constrained pose. Body inertia and joint stiffness naturally clean up minor noise and occlusions in the captured marker data. The resulting joint angles in transit allow the specification of the complete set of dynamic torques. To test this feature of HDM quantitatively, the recovered joints angles while walking according to the reduced marker set were compared with those from the full marker set. Fig. 9 illustrates the recovered joints angles are quite similar with the original joints angles.

**Fig 8.**
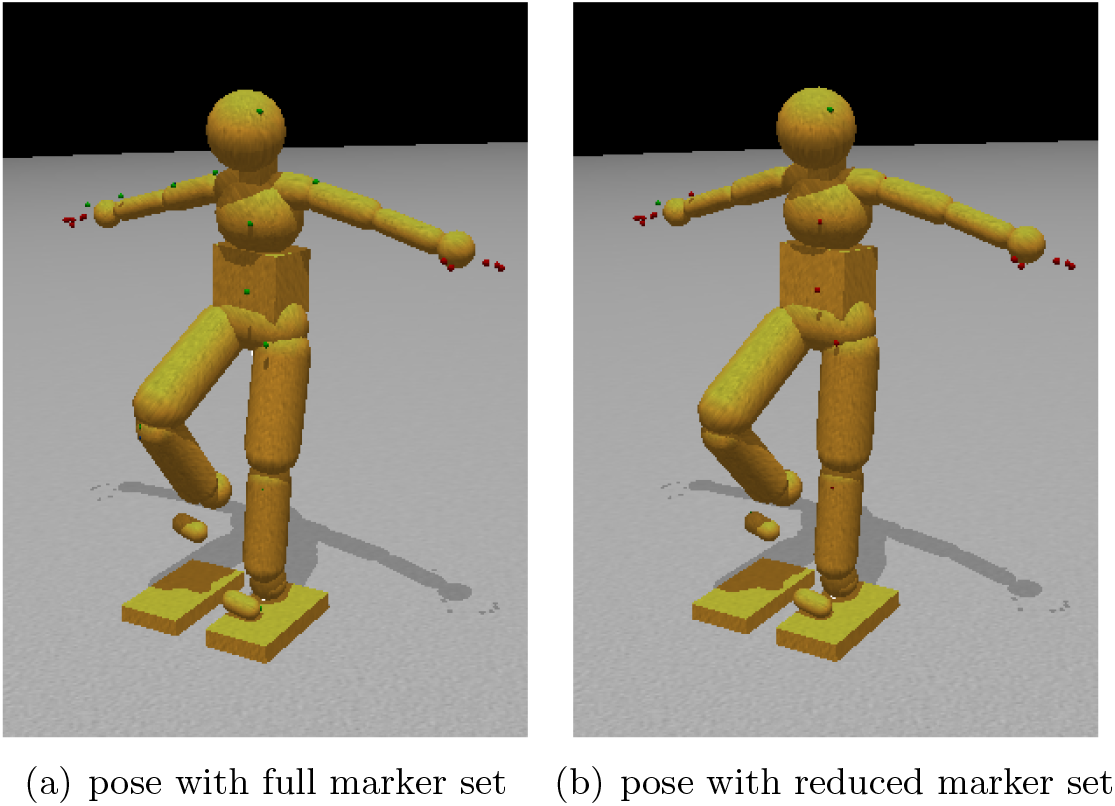
Movement control using dynamic synergies. (a) Body configuration using all marker constraints.Note the similarity to the sparsely constrained pose. (b) Body configuration using constraints on only the head, hands, and feet. In many cases, the pose found using a full set of marker constraints is quite close to that found by a sparse set of constraints. These two images show almost no differences between using a full or a sparse set of marker constraints.

**Fig 9.**
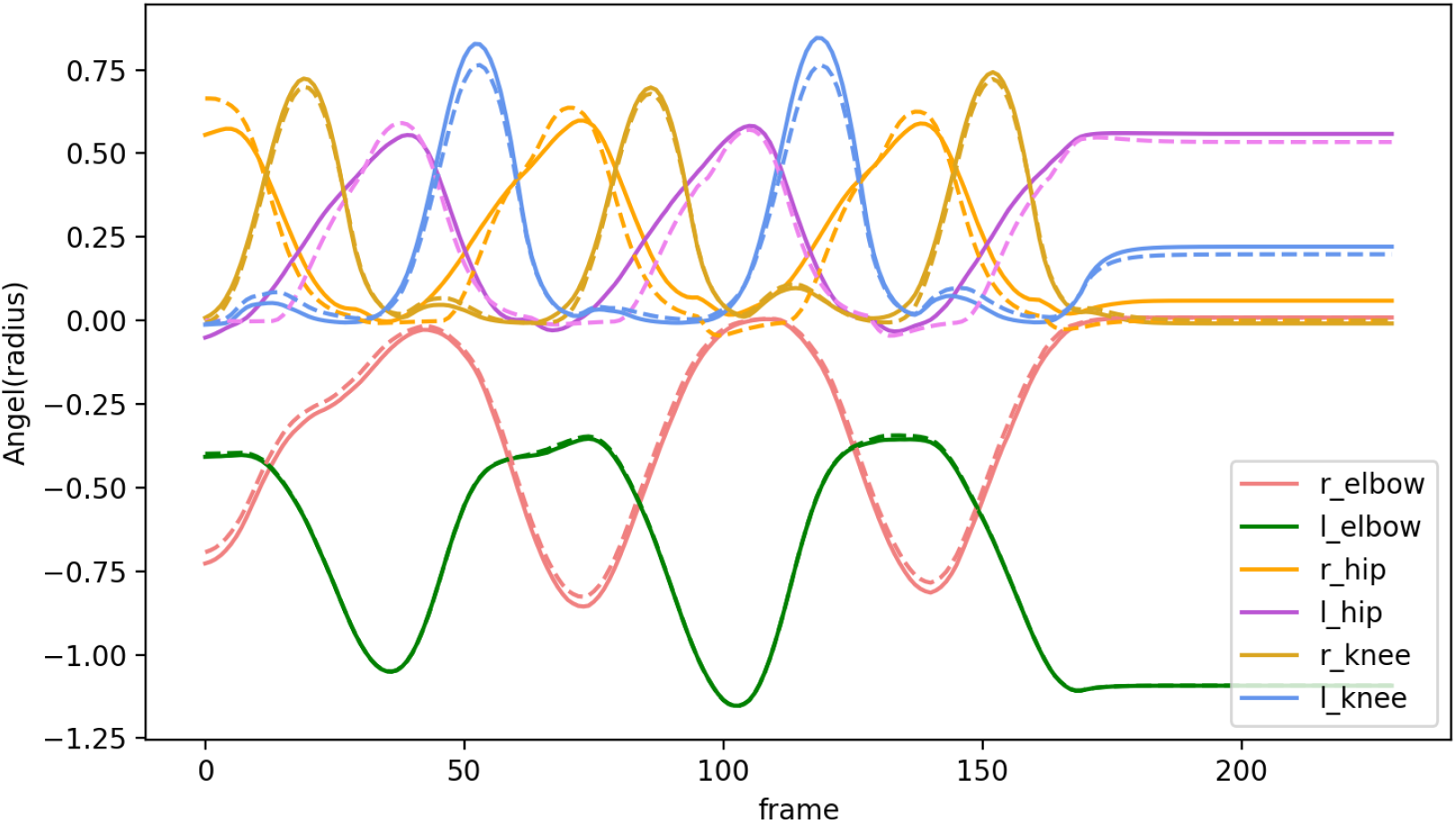
Comparison of joint angles along selected degree of freedom. Solid lines show joint angels recovered based on full marker set. Dashed lines show joint angels recovered based on reduced marker set.

This result has important general implications. First of all, the finding suggests that the kinematic plan for movements can be compressed into a subset of formative trajectories, leaving the remaining degrees of freedom interpolated using the body’s dynamic constraint. Another aspect of this observation is that the reduced set can be used to adjust movements to individual circumstances, again leaving the detailed interpolation to the dynamics.

## Discussion

The paper has aimed to publicize a novel system for quantitatively modeling whole-body movements. Its 48 degrees of freedom and generalized spring constraints allow models of scale that are robust to disturbances. In addition to being an analytical tool, it can also generate movements from a kinematic plan.

The core of our simulations exploits the observation that realizations of constraints behave like implicit springs. The parameters that soften constraints into springs exhibit many advantageous properties. They stabilize the simulation, pushing a constrained system away from singularities, and reducing constraint error. A fundamental question concerned with the HDM system is whether it can recover the trajectories and the cost of a known physical system’s complex motion. The experiments described above showing the HDM synthesizing human motions with high levels of accuracy. One further principle behind our tests is that one way of illustrating the method’s robustness is to combine a kinematic data set from the source with another set of dynamic parameters. In tests, the data gathered with a different motion capture device is combined with the inertial data from another model to make a composite. Our tests used the Carnegie Mellon University’s graphics laboratory’s motion capture database ^5^. This beneficial and extensive database contains whole-body motion data sets for different human subjects performing various natural motions. The database was created by motion capture, and the positions of markers on the bodies are one of the primary sources of motion data. We did not know the individual dynamic parameters. However, by adopting the database’s marker conventions, we could use our dynamics calculation to compute joint torques for the hybrid system. Although the estimate is thus done for a synthetic pairing of kinematic data and dynamic parameters, the point is to show that, even with this combination, the integration is stable and leads to identifiable torques.

A central feature of the system is the production of the movements’ energetic cost to provide the capability to compare different movement scenarios. Achieving this aim can be tricky, owing to the lack of systems that can provide independent cost measures. Energetic cost measurement of human movements has been studied for decades. The most straightforward and frequently used method is to measure the metabolic cost,e.g., subjects breath through a mouthpiece to measure rates of oxygen consumption (VO2) [25–31]. Measuring the changes in muscle coactivation and stiffness using Electromyographic (EMG) is another common way to reflect metabolic changes [32]. However, these methods are time-consuming, and the required configuration restricts the variety of experiments. For example, the VO2 process does not work for virtual-reality tasks as subjects need to wear the VR helmet on their head, leaving little space for a mouthpiece.

By comparison with the above methods, the HDM provides a stable and versatile platform with several uses. One is the use with force plates, as shown in our experiment, to measure the stance’s change. Another option is to use the HDM system to produce correlations with similar tests with human subjects, such as our research with stiff-arm walking. Once we have vetted the system in many such areas, it can be used as a predictive tool, as in the experiment showing the different costs of reaching targets. We have developed a large-scale three-dimensional tracing experiment in virtual reality [33] to elicit natural whole-body movements under common goals. Our future work is to analyze the energetic cost using the HDM.

Besides its use of a mechanism for interpreting experiments, the system can also serve as a good base for theorizing about the human system’s organization concerning its space-time performance since many of these issues are open. While an enormous amount of research in human motor control has produced ever more refined subsystem components’ elucidations, a comprehensive theory at the level of large scale dynamics is still unsettled. One main obstacle is a description of how the motor cortex can communicate control information to drive the high temporal bandwidths of the spinal cord circuitry. Several possibilities were debated at the *Neural Control of Movement* conference in 2013 without definitive result. We have emphasized is that the motor cortex communicates a coded kinematic plan together with stiffness settings. A study with kinematics coded with temporal basis functions has shown that a kinematic plan can be coded to reduce the bandwidth needed by a factor of approximately 10^3^ [34,35]. The HDM shows that such a model can play a useful role in studying the kinematic-plan model’s consequences. In particular, the reduced degree of freedom control demonstration supports the *uncontrolled manifold* view wherein a subset of crucial degrees of freedom can direct a movement with the uncontrolled degrees of freedom interpolating the movement using the system’s dynamics [36, 37]

In regard to the uncontrolled manifold concept, a very important insight was the use of reduced degrees of freedom constraints in computing the dynamics. If the limitations are near the number of DOFs of the system, then the torque recovery can quickly become numerically unstable. However, between 20 to 41 markers in the HDM provide sufficient constraints to integrate the dynamic equations reliably by allowing the system’s natural dynamics to interpolate the motion appropriately.

The method has several advantages over alternative methods. First, it can be easily implemented in a single robust framework of the physics engine. Using the physics engine for multiple tasks allows a unique human model to be used from start to finish, rather than being forced to use the conventions built into a commercial package. Second, the method is fast. The simulation engine is designed for performance, making it possible to analyze movement in real-time and create interactive experiments with stimuli dependent on the feedback results. Third, the software is free. Freely accessible code, such as ODE, is useful because it facilitates comparison and collaboration in research. Fourth, the method handles multiple ground contacts and noisy data challenging to related approaches. Kinematic loops do not require any special treatment. The method is robust even to large perturbations making data dynamically inconsistent. Finally, the tunable parameters (CFM), couched in the physics framework, are intuitive. It is more straightforward to specify the importance of a constraint in force and mass rather than arbitrary gains and weightings. We illustrate these advantages by using ODE to analyze and reproduce movement recorded from optical motion capture.

There are several ways to improve the system, but three are the most important. One limitation of our method for computing torque is that it is insensitive to muscle stiffness, which is both passive and can be actively modulated [38, 39]. Increasing stiffness will increase the overall net movement energetic cost and needs to be taken into account. The observation somewhat ameliorates this issue that in most natural tasks, subjects will try to minimize energetic costs and thus exploit natural dynamics whenever they can [36, 40, 41], reducing high levels of co-contraction. However, the ubiquitous use of spring as constraints means opening up the possibility that one can add springs to the joint degrees of freedom to model stiffness. These could also have parametric programmable spring constants to model muscle co-contraction. The second feature that could be added is a system to keep the human model upright. Any of the three human sources of this needed information - visual, vestibular, and proprioception - would be candidates for this practical constraint. At present, the HDM uses a faux system of rotational torques at the center of gravity, but these could easily be replaced with more appropriate ankle torques. The third feature to be added is the separation of gravitational torques from control torques as only the latter effect metabolic cost directly. This improvement is a matter of modifying ODE’s low-level code, and the plan is that this will be tackled shortly.

In summary, the forty-eight degree of freedom dynamic human model is a fast, robust, intuitive, and inexpensive multi-purpose tool for simulating, analyzing, and synthesizing humanoid movement. The system’s capability is a very stable set of integrations that readily handle the inclusion of multiple points of surface contact. The HDM uses a closed-loop step at each time step so that the computed torques are appropriate for the new posture. In contrast, when the computed torques are saved and replayed, small errors in the kinematics accumulate. Each set of torques is no longer appropriate for the computed posture, and the overall system rapidly becomes unstable. These results’ significance extends beyond the simulation stability issue and provides a strong argument for the suitability of the kinematic plan’s close-loop control as a biological model.

## Methods

A novel way to compute the energy cost of human movements has been developed by building a human dynamic model on the top of a physical engine ODE.

### Human Model

Our techniques use a simulated model of the human whose movement is analyzed. The first order of business is to build a physical model capable of representing human movements, of which the accuracy influences the outcome of the analysis. Fig. 10 shows the body segments and topology of the model. The humanoid model is a collection of rigid bodies connected by joints. Each joint connects two rigid bodies with anchor points (center of rotation) defined in the reference frame of both bodies. The locations of these anchor points determine the segment dimensions (bone lengths) of the character model.

**Fig 10.**
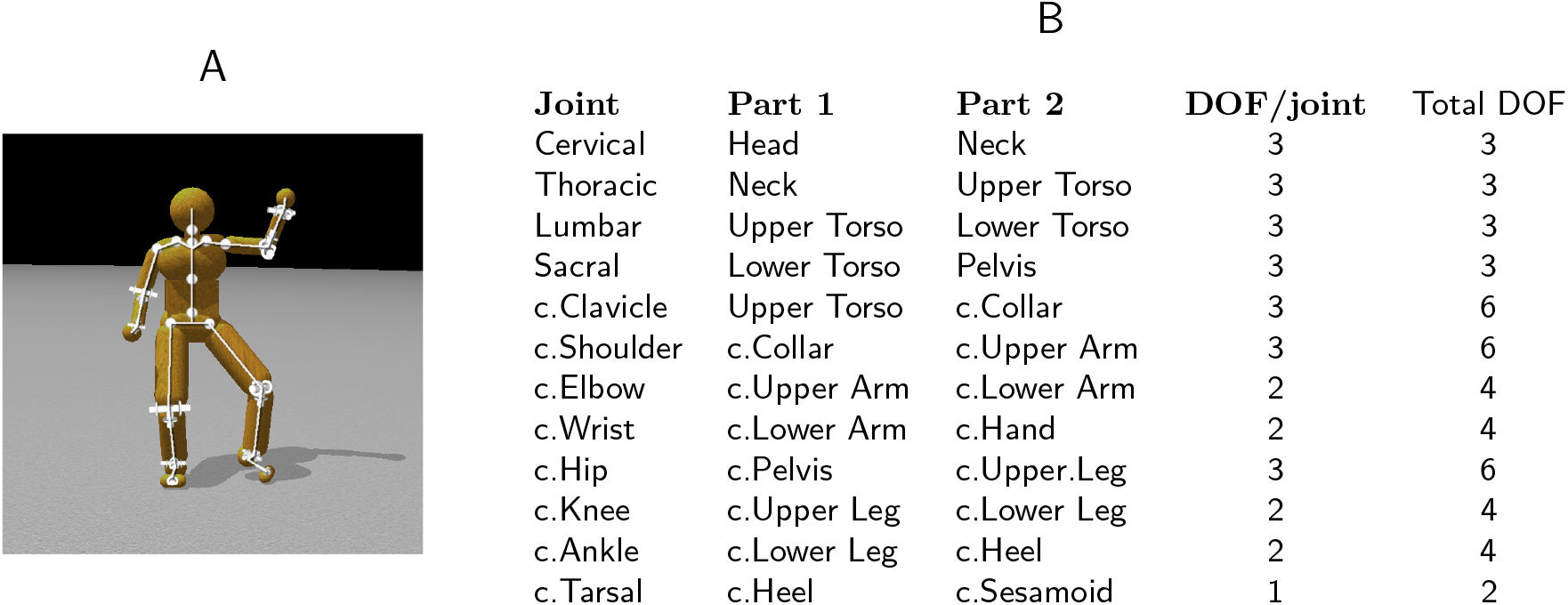
The 48 degree of freedom model. A. Four ball-and-socket joints connect five body-segments along the spine from the head to the waist. Ball-and-socket joints are also used at the collar-bone, shoulder, and hip. B. A summary of the joints used in the model. c. = chiral: there are two of each of these joints (left and right). Universal joints are used at the elbows, wrists, knees, and ankles. Hinge joints connect the toes to the heels. All joints limit the range of motion to angles plausible for human movement. Our model assumes that joint DOFs summarize the effects of component muscles.

#### Model degree of freedom details

The model structure consists of 21 separate rigid bodies connected by 20 joints (Fig 10). The relative orientation of some bodies is constrained by using universal joints for the elbows, wrists, knees, and ankles and hinge joints to connect the toes to the heels. Universal joints restrict one angular degree of freedom; e.g., when the arm is bent at the elbow, the forearm cannot rotate around the principal axis of the upper arm unless the upper arm itself rotates. However, the forearm can rotate at the elbow around its own principal axis (modeling the twisting movement of the radius and ulnar bones). All other joints are left as ball-and-socket joints with three angular degrees of freedom: hips, shoulders, collar-bones, upper-neck, lower-neck, upper spine, and lower spine. This arrangement of joints leaves a total of 48 unconstrained internal degrees of freedom. An advantage of building humanoid model in this way is that joint connections are not treated as holonomic (perfectly rigid) constraints, but rather as very stiff springs that hold body parts together like tendons and muscles.

### Data Fitting

The technique for fitting a model to data begins with a character model that serves as a template, Fig. 10, providing the number of body segments and topology of the model. We further require that labeled markers used in motion capture be assigned to specific model segments. It may be straightforward to derive these using a technique such as in [42, 43]. However, it is also not difficult to do by hand. It would become tedious if one had to go through the process for many different models. Fortunately, the motion capture suit typically puts the markers on the same body segments (Fig. 11), even if they are in slightly different places, and the body segments have different dimensions.

**Fig 11.**
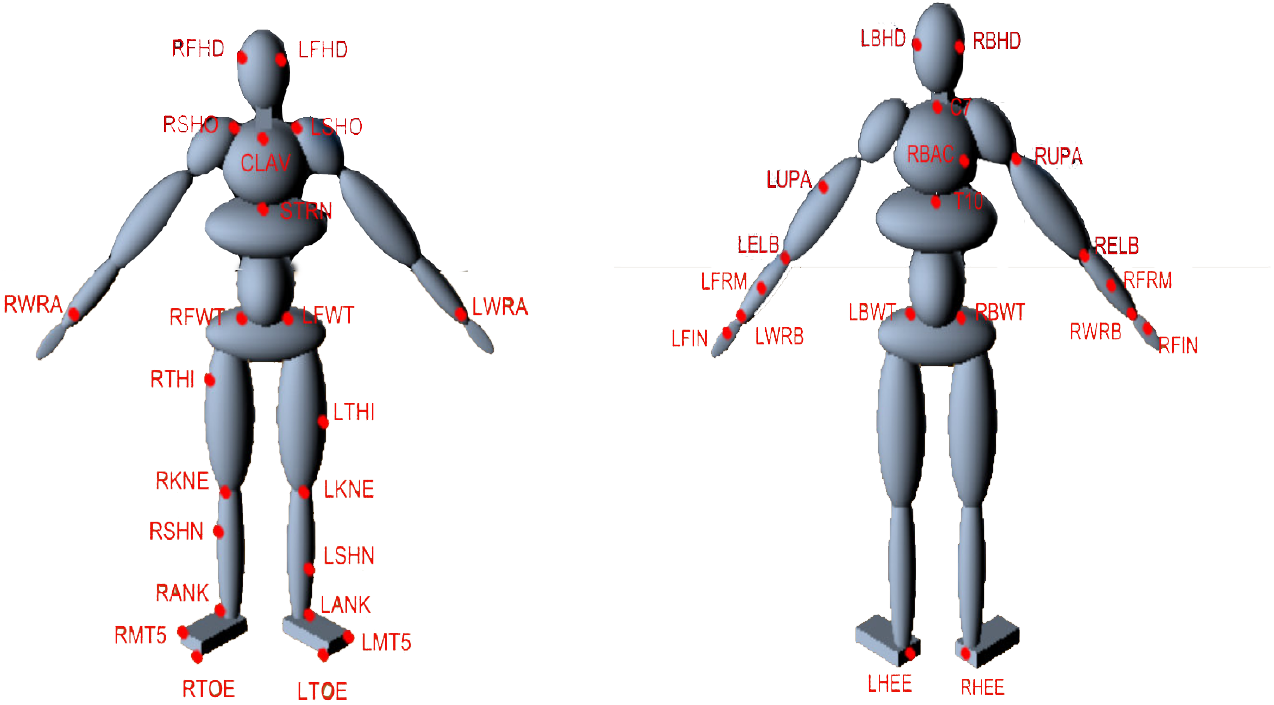
Marker arrangement on the motion capture suit. The suit contains 51 markers as shown by the LEDs in total but only 41 are used in the model e.g. Markers that are used are present on the fingers. Markers can easily assigned to specific model segments. For example, the markers of RBHD, RFHD, LFHD and LBHD are assigned to Head segment while the markers of RBWT, RFWT, LFWT and LBWT belong to Pelvis segment.

We present a method in S2 Appendix section, for using marker data to help determine the dimensions of the model segments and where markers attach to the model. Although this method could easily be automated, in practice, the research did not rely on very many different models and so the system uses a mechanism for relaxing the marker attachment points and joint anchors with the click of a button in the graphical user interface (Fig. 1). With a new data set, a handful of iterations proved sufficient to produce a reasonable model with marker attachments that fit the data well enough for further analysis. This algorithm does not address joint limits on a range of motion. These can also be learned [44], but in our case, the range of motion for each joint is set *a priori*. After determining segment lengths, we set other segment dimensions as appropriate to fit against the markers. Mass properties for each segment assume uniform density by volume.

Given motion capture data of a subject, the model is fit to the subject’s dimensions and joint-range-of-motion is constrained to approximate the subject’s flexibility. Additionally, the model segments have inertial matrix properties. The initial mass assignment to each segment assumes a uniform density of water 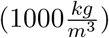 for the volume associated with each rigid body. The mass assignment should be modified to roughly match that of a specific subject. The increased fidelity, required for individual subjects in clinical biomechanics research would employ more sophisticated techniques for a better approximation of mass distribution in the model. Interestingly, however, the experimental results discussed above show that even this low fidelity model is sufficient to produce high-quality data that compares favorably with data gathered from independent sensors.

### Pose Fitting

Having addressed the issues in attaching the model to motion capture data, we turn to the construction of its capability of representing human movements. Various commercial packages provide different methods for converting marker trajectories into sequences of body poses, but they can be time-consuming, expensive, or difficult to use. This section describes an approach related to [45] and [46] that is free, fast, uses intuitive parameters, and allows the user to fit markers to whatever model they wish.

The method uses the physics engine to constrain a character model to fit marker data and other constraints. Markers are modelled as infinitely massed points attached to the character model. Given a frame of marker data, the position and orientation of all body segments can be found by balancing internal joint targets and external marker data. From the global position and orientation of the different body segments, it becomes a simple matter to compute relative orientations (joint angles).

The internal degrees of freedom are limited by range of motion constraints, e.g. the elbows and knees cannot bend backwards. All other joints have similar range-of-motion limitations based on the subject’s flexibility. Furthermore, each joint is set to have a “target state”, a preferred relative orientation between its connected bodies. These preferences can be thought of as “muscle stiffnesses” and are modeled as weak constraints with limited force. Joint limits and stiffness serve as a prior over possible poses so that in the absence of any marker data at all, the model still takes on a pose. Consequently, every internal degree of freedom is constrained to some degree. These constraints hold the model together and give it a default pose. Marker data pull the model from the default pose into a new pose, e.g. Fig 12. For a given frame of motion capture data, each marker is connected to a body segment using a ball-and-socket joint constraint. A total of 41 markers, which do not contribute any degrees of freedom because of their infinite mass, attach to the character model, adding an additional 3 × 41 = 123 constraint dimensions.

**Fig 12.**
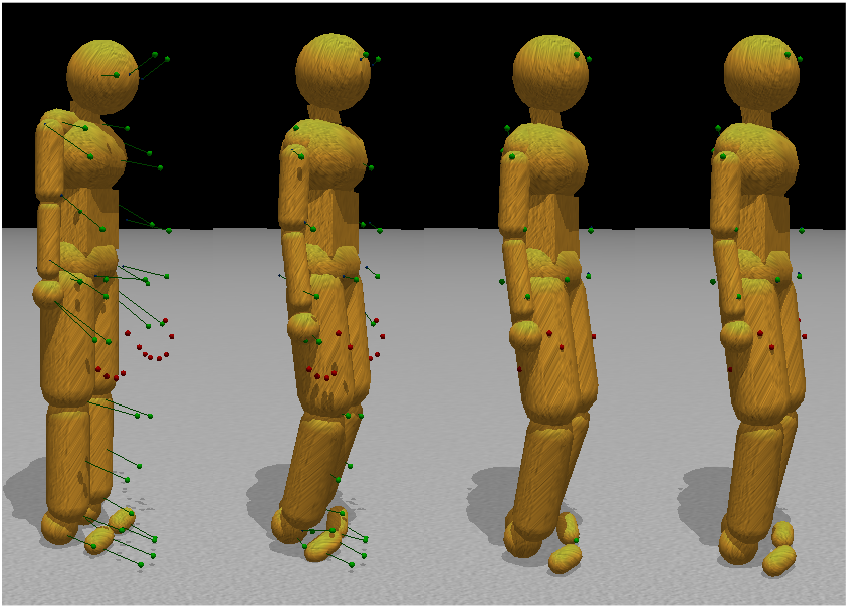
Pose fitting. Initially the motion capture data points are in a very different configuration than the initial stance of the model. To find the appropriate correspondences, simulated markers attach to the humanoid model through ball-and-socket joints and pull the body parts into place, subject to model joint constraints. The left to right sequence in the figure shows the body targets being gradually reconciled with the external markers.

Finally, collisions between the ground and the feet also influence the model pose. Each foot can form up to three contact points with the ground. Inequality constraints at these points prevent penetration with the ground. When both feet are firmly on the ground, all markers are actively pulling the body into a pose, all joints are holding the body together, and joint limits and stiffness are biasing the relative orientation of the bodies. The experiments described above show that the model can simulated the ground force correctly.

This approach is simple intuitive: attach markers to the model with springs and then drag the body along. The parameter, tunable for each constraint, which is known in ODE as the constraint force mixing parameter (CFM), allows a constraint to slip proportional to the amount of force that would be required to maintain the constraint. For the regular internal body joints and contact constraints, we use a CFM value of 1×10^*−*5^ while for the constraints between markers and body parts we use 1×10^*−*4^. Both of these values represent very stiff springs although they are different by an order of magnitude. This stiffness stabilizes the simulation by allowing the markers to stretch slightly from their mapped locations in the event that the marker constraints are not compatible with the character model. Fig 12 shows that when the markers move, the constraints drag the character along with them.

### Inverse Dynamics

It can be useful to know the torques to apply at each joint or the required effort to accomplish a particular movement. Given a kinematic sequence of body poses, the physics engine ODE can archive the computation with minimal effort. Given the constraints, such as each joint’s angular velocity, it can correctly compute the desired torques/forces measurements.

The process is straightforward. Given the current joint angle and the desired joint angle for the next frame, the relative angular velocity of the body parts is constrained as to achieve the target orientation on the next frame. Contact constraints are necessary to prevent ground surface penetration as well. The ODE physics library handles the constraints and solves the torques and forces that are used to satisfy each constraint in the process.

For computing inverse dynamics, the first step is to initialize the model to a starting dynamic state. The initial state can be found from the first and second frames of kinematic pose data. The model pose is set by using the second frame of data, and the initial linear and angular velocity of each joint is computed by taking the finite difference between the two frames (and dividing by the timestep). Computing velocity through finite differences is appropriate for a physics engine using first-order semi-implicit Euler integration. After that, continuously find the torques between two consecutive frames of pose data using the finite difference between poses to compute angular velocities that will move the model from the current to the next pose.

Differentiating again, this time between the current and future velocity gives a target acceleration that becomes a constraint on the model. The primary difference between this step and the previously discussed method for finding pose from marker data is that there are no marker constraints dragging the body into place, and the internal muscle stiffness drives the model toward a target pose on each frame instead of toward a ‘default’ pose. Because there are fewer constraints in play, stiffer muscle forces are used, but the absolute forces the muscles can apply are limited to prevent muscle forces from being unreasonably large. Again, in this case, we can use the relative spring stiffnesses to express the confidence in the measurements. We use very stiff springs (CFM = 10^*−*10^) to keep the model segments together. We use looser constraints to keep the feet from penetrating the ground (CFM = 10^*−*5^) and to constrain the model to adopt the appropriate pose (CFM = 10^*−*8^)

#### Residual torques/forces

The torque calculation by the HDM is ideal in the sense of solving the dynamic equations, but in the actual situation there needs to be a corrective system for incidental errors. In the human system there are multiple corrective system based on vision, proprioception and the vestibular system. Such corrective systems have been extensively studied e.g. [18, 19, 47].

In classical inverse dynamic area, discrepancies between the model and human that created the data necessitate non-realistic “residual” forces to keep the model from falling over when dynamically reproducing most movements. A 6 degree of freedom joint between the waist segment and the global frame generate the external forces. A weak, limited spring constrains the waist segment to achieve its recorded pose relative to the global frame. The experiments show that attaching the external constraint to the head or the feet has little noticeable difference. The non-realistic external forces (residuals) account for noise as well as discrepancies between the model and the human generating the data. In particular, differences in how the feet interact with the ground cause errors in our analysis. In most cases it is only necessary to constrain two of the 6 angular degrees of freedom (pitch and roll), leaving the other four external degrees of freedom disabled. The two angular constraints keep the body from falling over but allow it to move about through simulated ground interactions.

The stabilization system completes the model. It can be implemented in parallel, with the control used to stabilize the residual necessary to balance. With this included, The simulation can reproduce highly dynamic motions, e.g. see Fig 2, which shows a jumping sequence made originally by a human subject and recreated using the torques computed by the inverse dynamics model.

### Method summary

For each human subject we construct a dynamic model and force that model to follow the subject’s motion capture data, which leads directly to the recovery of joint angles. Our algorithm constrains the dynamic model to track these angles and consequently can estimate the correct joint torques. This concept was originally demonstrated in two dimensions for human walking by [48]. We have extended the method to the significantly more demanding case of 48 DOFs in three dimensions and arbitrary posture changes. Fig. 10 lists the body segments. The dimensions of each segment are matched those of an individual subject. The principal difficulty is that the constraints in the high DOF 3D model present many delicate numerical issues for the ODE solver that need to be addressed [14]. Currently the dynamic model does not attempt to model stiffness components, with the consequence that it can only directly recover the net torques at each DOF.

The body segments are used by the simulation for both collision detection and the calculation of mass properties. Mass and inertial properties are computed from the volume of the body parts using a constant density of 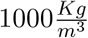. The dimensions and articulation are designed to allow the model to reproduce most movements the human can make. For example, joints at the elbows have two DOFs to reproduce the hinge movement of the elbow as well as the twisting movement of the radius and ulna bones in the arm. Joint DOFs are also limited to prevent impossible movements such as reverse bending of the elbows or knees.

For data capture a subject wears the motion capture system developed by PhaseSpace. Each PhaseSpace LED marker is mapped to a corresponding point on the model. The markers are then introduced into the physics simulation as kinematic bodies without collision geometry. As a heuristic, each marker kinematic body is effectively treated as having infinite mass so that when another dynamic body is attached through a joint constraint to a marker, only the dynamic body’s trajectory can be changed by the constraint.

The PhaseSpace motion capture system records 41 3-dimension positions of specific human body locations over time and maps these markers to appropriate locations on the model. When the simulation is stepped forward, a constraint solver attempts to find a body state that satisfies the internal joint constraints, the external marker constraints, and other constraints such as ground forces, joint stiffnesses, and conservation of momentum. Knowing the kinematics allows the recovery of the dynamics, since the joint velocities allow the equations of motion to be inverted. The retrieved forces can be used to generate feed-forward torque profiles for actuating the character.

The overall idea behind the method for calculating joint torques is straightforward and has been implemented in ODE. The mathematics underlying the rigid body simulation software used in our work is explained in the S1 Appendix section.

## Supporting information

### S1 Appendix

The principal insight in this section is that ODE can be used as an effective controller. We present a derivation of the mathematics underlying the physics simulation. The derivation comes from directly analyzing the ODE codebase and it consequently differs from other derivations using Lagrange multipliers to arrive at the same final result (e.g., [49]). We present another derivation illustrating the equivalence between softened constraints in ODE and implicit springs.

When modeling human movements, we assume that the human body does not collide significantly with itself and so typically only process collisions between the model and the ground. Collisions between the model and the ground, however, play an important role in analyzing and synthesizing movement data such as walking. Collision handling involves creating a constraint between the colliding bodies and is the primary contribution of the model. We will describe this methodology after first introducing the physical simulation details.

#### Notation

Physical simulation involves a large number of different variables to represent constraints and relevant physical quantities. Table 1 presents specific symbols and their meanings for reference.

**Table 1.** Meanings of specific symbols used to discuss dynamic simulation

| Symbol | Meaning |
| --- | --- |
| $\mathbf{x}$ | position or state of one or more rigid bodies |
| $\dot{\mathbf{x}}$ | velocity (usu. linear and angular) |
| $\ddot{\mathbf{x}}$ | acceleration (usu. linear and angular) |
| $\mathbf{R}$ | rotation matrix representing orientation of a body |
| $\boldsymbol{\omega}$ | angular velocity |
| $\mathbf{q}$ | quaternion representation of an orientation or rotation |
| $m$ | mass of a single rigid body |
| $\mathbf{M}$ | mass matrix |
| $\mathbf{I}$ | identity matrix |
| $\mathcal{I}$ | moment of inertia tensor |
| $n_b, n_c$ | number of bodies, number of constraints |
| $\boldsymbol{\alpha}, \boldsymbol{\beta}$ | stabilizing parameters added to the equations of motion |
| $\phi()$ | error or energy function for a single constraint |
| $\mathbf{J}$ | matrix of partial derivatives of constraint error functions |
| $h$ | timestep |
| $\mathbf{f}$ | forces (and torques) |
| $\boldsymbol{\tau}$ | torques |
| $\boldsymbol{\lambda}$ | constraint forces |

Scalars are represented with lower-case, un-bolded symbols: *x*. Bold lower-case symbols represent column vectors,

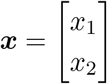

Bold, upper-case symbols to represent matrices:

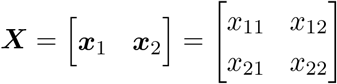

Dot-notation to indicates time derivative: 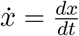. The circumflex accent indicates a 3d vector being used as a skew-symmetric matrix representing a cross-product operation:

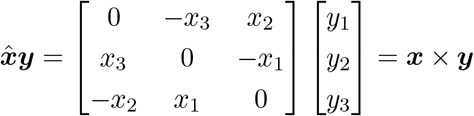

Coordinates are typically relative to a global reference frame. However, a tilde, 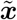, indicates a quantity that uses a local reference frame, e.g., a body-relative frame, rather than the global frame. We use subscripts to indicate that a quantity refers to a specific dimension, a particular rigid body, a point in time, but clarify the subscript’s meaning when necessary to remove ambiguity. Table 1 introduces the primary symbols within the text.

For conciseness in notation, we typically combine angular and linear quantities as a single symbol. This representation is used both for position and orientation even though orientation does not conveniently fit into a 3×1 vector. Fortunately, angular velocity and angular acceleration, ***ω*** and 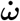, do combine well with linear velocity and acceleration 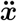 and 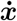, and it is these quantities that feature primarily when dealing with a constrained system. We will also represent the state of multiple bodies using a single symbol when convenient. For example, for a system with two bodies, we will represent the combined linear and angular accelerations (a 12d vector) as 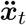. For this same 2-body system, Newton’s law relating force, mass, and acceleration is as follows:

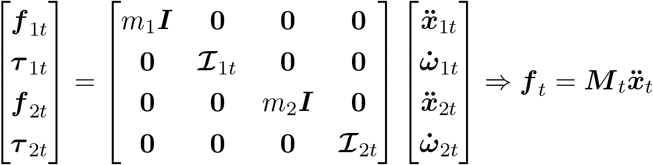

where ***I*** and 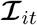 are 3 × 3 block matrices.

#### Dynamic State

Coordinates in the simulation world are defined relative to an arbitrary origin and basis set of directions. We refer to this inertial frame as the “global frame”. Each rigid body also has its own point of reference and set of directions. Any point in the global frame can also be described relative to a body’s frame of reference. It is convenient to define the point of reference of a body as its center of mass and use its principal inertial axes of symmetry as directions.

The position of the center of mass and orientation of a body within the global frame are here defined as ***x*** and ***R*** respectively. In 3d space, ***x*** is a 3 × 1 vector:

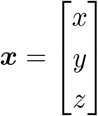

where *x*, *y*, and *z* are the distance from the origin along each of the three directions that establish the global frame of reference. For consistency, we deal with these distances in meters and assign “up” to the positive *z* axis. The orientation ***R*** of a body is a 3 × 3 orthonormal matrix whose columns give the body’s local direction frame relative to the global frame:

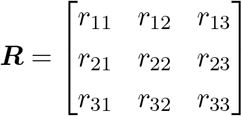

Conservation of momentum makes it necessary to keep track of the time derivative of these quantities: 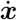 and 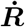. Instead of explicitly representing 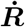, it is convenient to keep track of the angular velocity:

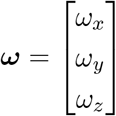

The relationship between these quantities is

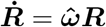

Representing orientation as a 3 × 3 matrix can be unwieldy. To properly represent an orientation (or pure rotation) it must be an orthonormal matrix. An orthonormal matrix uses nine elements to represent a property with only three degrees of freedom. Unfortunately, any three-element representation of orientation suffers from singularities [50]. We make use of unit-length quaternions to represent orientations and changes in orientation. Quaternions are convenient because of their close relationship to angular velocities. Quaternions are similar to an axis-angle representation of a rotation. A quaternion ***q*** represents a rotation by *θ* around unit vector ***v*** with four elements:

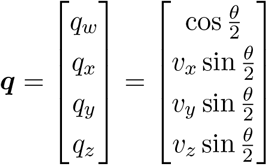

From an arbitrary angular velocity ***ω***, we can make a quaternion that represents the change in rotation that would occur during a timestep of *h*. After finding the amount of rotation *θ_t_* = *h ||**ω**_t_*||, one might naively find a “rotation quaternion” by normalizing ***ω_t_*** and then re-scaling by 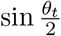 for a final quaternion: 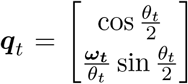. However, the normalization step becomes unstable as *θ_t_* approaches zero. To avoid that instability, we use the “sinc” function where 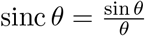. The sinc function allows us to remove the discontinuity that would result from division by zero and adds numerical stability. When *θ* is small, sinc (*θ*) can be approximated to within machine precision using the first two non-zero terms of its Taylor expansion (see [50]). The result is a discrete-time “rotation quaternion”:

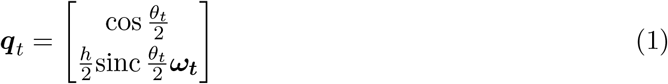

Given *n_b_* bodies, the dynamic state of the *i*^th^ body at time *t* is its position, orientation, linear velocity, and angular velocity: 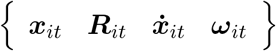. We will assume that all of these values are framed in the global coordinate system unless specified otherwise. The body dynamics are also affected by the body’s constant mass *m_i_* and inertia tensor 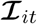. The moment of inertia tensor, 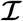, is indexed by time because the body’s orientation changes how the the mass is distributed relative to the world frame: 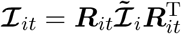. We assume that the inertia tensor is constant relative to the body-local frame of reference (i.e., bodies are rigid).

In simulation, the forces ***f*** applied to the rigid bodies come from three general sources. These are constraint forces (***f_c_***), gravitational and gyroscopic forces (***f_g_***), and user/control forces (***f_u_***): *f* = *f_c_* + *f_g_* + *f_u_*.

#### Integration Step

When a force is applied to a body, it translates into acceleration that is inversely proportional to the mass. Velocity is the time integral of acceleration, 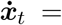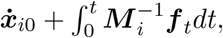, and position is the time integral of velocity, 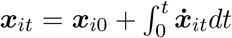. Because ***f** _t_* may depend on ***x**_t_* and 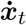 as well as on discontinuous collisions and control inputs, analytic descriptions of body state are not usually possible. Instead we discretize the equations of motion and use a small, discrete timestep, *h*, to numerically approximate system dynamics. The most obvious thing to do is to linearize the force function, ***f** _t_*, and then take all the quantities from time *t* and use them to find the state at time *t* + *h*:

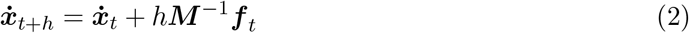

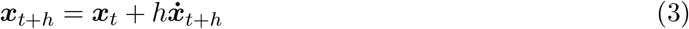

This “semi-implicit Euler” integration uses using the future velocity for computing position and is more stable than the standard formulation.

Although we lump orientation and position together as a single symbol, in practice there are a few distinctions that need mentioning. For example, gravity only applies to the linear state, while gyroscopic torques only apply to angular state. Gravitational forces are very straightforward, ***f***_grav_ = ***Mg***, where ***g*** indicates the direction and magnitude of gravitational acceleration and is often very simple; e.g., for a single rigid body ***g*** = [0 0 −9.8 0 0 0]^T^.

Rotation is a non-linear phenomenon. However, we can approximate the motion of a rotating body by adding torques that imitate gyroscopic effects, see [51]. Gyroscopic torques are applied to maintain conservation of angular momentum. Explicitly applying gyroscopic torques to bodies allows us to treat the rest of the system as though it conserved angular velocity rather than angular momentum. Thereafter, we can deal with the combined linear and angular quantities as a linear system.

The gyroscopic torques for each body are linearly approximated by

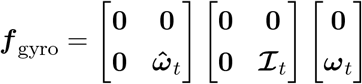

These forces are zero if the three principal moments of the inertia tensor are equal. Otherwise, they represent the forces necessary for conservation of angular momentum. Unfortunately, this approximation tends to introduce energy into the system. We have reduced this problem in ODE by adding in additional terms as described in [51].

The constrained system is solved using mostly accelerations and velocities. At the end, however, it is necessary to integrate the velocities into new positions and orientations. Position and orientation are updated differently. For position, it is sufficient to multiply the linear velocity by the timestep and add it to the current position. Adding angular velocity to orientation is not as straightforward. We integrate angular velocity into orientation by converting ***ω***_*i*(*t*+*h*)_ into a quaternion and then use the quaternion to rotate the current orientation forward in time following [50].

#### Constraint Equation

When a rigid body is moving or spinning freely through space, the integration equations are sufficient to simulate dynamics. Adding constraints modifies the bodies’ movements. Maintaining a relationship between two bodies requires forming a constraint on the state of the bodies. The integration equations tell us how to go from force to velocity and from there to position and orientation. To simulate an articulated model using maximal coordinates, we need to know what forces constraints apply to the bodies in the system.

To find the constraint forces, one must be able to mathematically describe the constraint. We define a multi-dimensional function over the combined position and orientation of all bodies in the system, ***ϕ***(***x**_t_*), that produces a vector of size *n_c_* specifying how much each constraint is violated, where *n_c_* is the number of constraints acting on the system. For example, if the *i*^th^ constraint keeps body *b*_2_ a distance *d* above body *b*_1_ in the *z* direction, we would have *ϕ_i_*(***x***) = *x*_2*z*_ − *x*_1*z*_ − *d*. If *b*_2_ is not separated from *b*_1_ by a distance of *d* in the *z* direction, *ϕ_i_*(***x***) reports the signed magnitude of that constraint error. For additional information on forming constraint equations, see [49, 52].

In general, the error for a constraint is non-zero. Given a measure of the error for a given state, we seek to find constraint forces, ***f_c_***, that reduce the error over subsequent time steps [53]. Specifically, over the timestep *h*, we seek a force to reduce the magnitude of the constraint error by a fraction *α*. That is

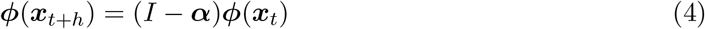

where ***α*** is a *k* × *k* diagonal matrix with each *α_i_* ∈ [0, 1] representing the fraction of error reduction over a time step. In ODE, the *α* value is controlled through the *error reduction parameter* (ERP) which can be set independently for each constrained degree of freedom. In practice, it is not possible to remove constraint error completely (*α* = 1) when using maximal coordinates because of error introduced by the various approximations employed to make the simulation linear and fast. Values of *α* typically fall within [0.2, 0.8]. Manipulating this value results in useful elastic and damping effects discussed later.

We use the symbol ***J**_t_* to represent the *n_c_* × 6*n_b_* matrix of partial derivatives of ***ϕ***(***x**_t_*). This matrix is a linear approximation of how the constraint error for each of the *n_c_* constraints changes when the positions and orientations of the bodies change:

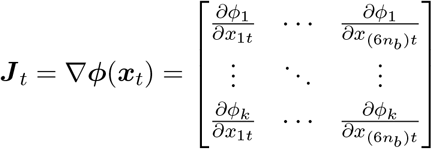

Finding the constraint forces that satisfy Eq. 4 involves removing all references to unknown future quantities. The Taylor expansion of ***ϕ***(***x**_t_*_+*h*_) at ***x**_t_*, truncated after the first order term, approximates the future constraint error:

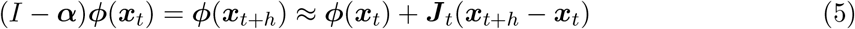

This truncation has the effect of treating all constraints as linear. Many constraints used to simulate various joints are linear; others, however, contain higher-order terms and this truncation is one potential source of error in simulation.

Combining the two integrator equations, Eqs. 2 and 3, gives the future position/orientation in terms of the present position, velocity, and forces:

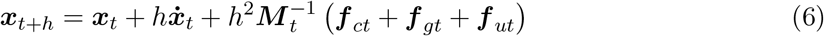

Equations 4, 5, and 6 combine to eliminate all references to future quantities:

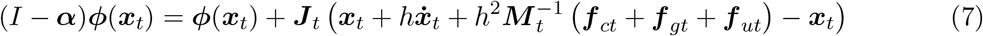

This leaves one unknown vector at time *t*: the constraint forces ***f** _ct_*. Rearranging and simplifying, we get

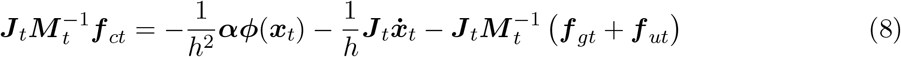

Note that in rearranging the terms this way, we divided both sides by the squared timestep, *h*^2^, effectively changing the problem from one dealing with positions to one dealing with accelerations. This conversion is possible because of the relationship established between acceleration and position by the semi-implicit Euler integrator.

Equation 8 is almost the equation that ODE solves when simulating physics. The right hand side is a desired acceleration. The first term on the right is the acceleration that would result in a velocity that would remove a fraction (*α*) of the constraint error. The second and third terms account for the effects of momentum (current velocity), gravity, and other forces (e.g., user control forces) applied to the bodies. Each constraint becomes its own dimension in a “constraint space”. The Jacobian matrix ***J*** projects accelerations from global forces into constraint space.

In general, the matrix on the left hand side of Eq. 8 is not square, making the problem under-constrained (or in some cases, potentially over-constrained). However, we can use d’Alembert’s principle [54] to restrict the constraint forces to lie in the constraint space.

Another method for arriving at the constraint equation is through the use of Lagrange multipliers. Consequently, the constraint-space forces are typically denoted with *λ*. The Jacobian transpose gives the relationship between a force applied in constraint space and force/torque applied in the full coordinate space: 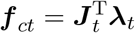.

The vector, ***λ**_t_*, holds the generalized forces applied by each constraint on all the bodies involved in that constraint, whereas ***f** _ct_* holds the sum of the constraint forces applied to each individual degree of freedom of each rigid body. The LHS of Eq. 8 can then be rewritten as 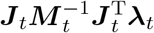, where 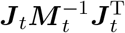 is now a *n_c_* × *n_c_* positive semi-definite matrix.

Returning to maximal coordinates, we will compress Eq. 8 down to

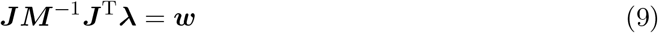

In general, the matrix ***JM** ^−^*^1^***J***^T^ may be singular. It is very easy to end up with redundant or conflicting constraints. For example, a box resting on the ground may get a contact constraint at each corner. If each contact prevents interpenetration and sliding (i.e., applies friction) then the contacts constrain a total of 12 degrees of freedom on a single rigid body with only 6 degrees of freedom to be constrained. Conflicting or redundant constraints can break the solver if not dealt with beforehand. The means for dealing with the conflict is clever. The physics engine softens the constraint, allowing it to “slip” proportional to the amount of force necessary to maintain it.

Because mass is always positive, the force, *λ*, applied to a particular constraint and the resulting constraint-space acceleration will have the same sign. Softening the constraint is therefore a matter of subtracting a scaled copy of *λ* from the desired acceleration (the right hand side): ***JM** ^−^*^1^***J***^T^***λ*** = ***w*** − ***βλ***, where ***β*** is an *n_c_* × *n_c_* diagonal matrix of (typically small) non-negative values. This subtraction, of course, is equivalent to adding ***β*** to the LHS. Adding these small values to the diagonal of the effective inverse-mass-matrix makes the constraints seem lighter to the solver and moves the matrix away from singularity:

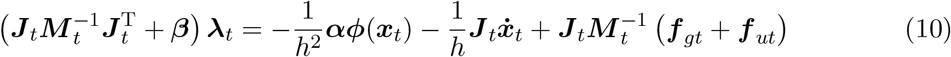

The original programmers built soft constraints into the ODE simulation code. The variable, *β*, tunable for each constraint, is known in ODE as the *constraint force mixing* parameter (CFM). At first glance, the addition of these parameters may seem loose and unprincipled. However, correctly setting the parameters, *α* and *β*, changes a hard constraint into a simulated implicit spring with first order integration (see [55]).

It is well-known that the formula for ideal damped spring force is identical to the formula for PD control. However, connecting these two facts, namely that (1) ODE’s constraints are mathematically equivalent to implicit damped springs and (2) damped springs are equivalent to PD controllers, has not been exploited. This insight is key to the success of the methods presented here. Our derivation shows that ODE’s constraints are, in fact, stable PD controllers along with examples of how to take advantage of this fact. We proceed by discussing proportional-derivative control and the mass-spring-damper equation.

#### Implicit Simulated Springs

Proportional-derivative (PD) control is a common method used to compute forces that drive a system toward a target state. The PD control equation is the same as a mass-spring-damper system. There are two parameters, *k_p_* and *k_d_*, that determine what force should be applied to a degree of freedom at any point in time. The stiffness, also called proportional gain (*k_p_*), specifies a force driving a degree of freedom toward its setpoint, 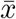 with strength proportional to the distance from the setpoint. The damping, also known as derivative gain (*k_d_*), counteracts the current velocity, slowing the system down to avoid overshooting. When a system uses PD control to encourage a degree of freedom to move toward a target state, the control force *f_ut_* at any instant in time is a function of the current position and velocity of the effective mass being controlled relative to its target:

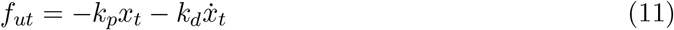

In a continuous time system, this controller is guaranteed to be stable as long as *k_d_* and *k_p_* are non-negative. With zero damping (*k_d_* = 0) the system oscillates in a sinusoidal wave pattern whose frequency is determined by the stiffness and mass and whose magnitude is determined by the initial conditions. With zero stiffness and positive damping, the velocity of the system decays exponentially with higher damping converging to zero more steeply. Discrete sampling of these forces, however, ruins the stability conditions. The potential for instability is apparent if we consider a mass *m* that only experiences damping forces. Using the semi-implicit Euler integrator, Eq. 2, we plug in the damping forces from Eq. 11 to get

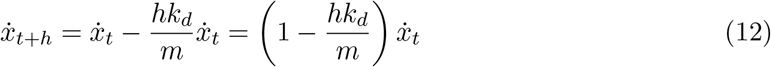

Time (*t*), mass (*m*), and damping (*k_d_*) should all be non-negative values. It is clear, then, from this equation, that if 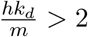, the velocity will oscillate between positive and negative values and grow in magnitude. This oscillation rapidly causes the simulation to “explode” and is annoyingly common when using PD control. Overly stiff springs hit a similar limit with explicit discrete integration that causes them to gain energy and explode. Consequently, explicit PD control gains are tricky to tune. They must fall within certain limits that depend on the timestep and the effective mass experienced by the system.

The cause for this instability lies in the discrete integration which is similar to approximating the area under a curve as the sum of multiple rectangles computed forward from the present (Fig. 13). One solution is to solve for the forces *implicitly*. Implicit integration is similar to approximating the area under a curve with fixed-width rectangles that end rather than begin on the curve. Rather than overestimate, this method tends to underestimate the area under an exponential curve. The resulting system does not explode, although it tends to dissipate rather than conserve energy. The implicit form of the damped-spring-law depends on the integrator it is applied to. Being ‘implicit’, in this case, specifies that spring forces are computed from the *future* state of the system. Consequently, Eq. 11 becomes the following, (note the changed temporal indices):

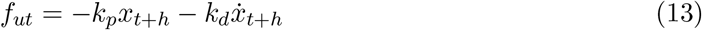

We do not know the future position or velocity, but using the integrator equations, Eqs. 3 and 2, we reframe Eq. 13 in terms of the current quantities and then solve for *f_ut_* to get

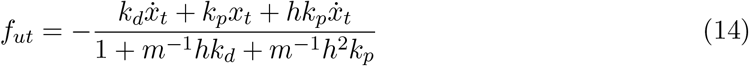

If we analyze a pure damped system as before but using Eq. 14, we end up with

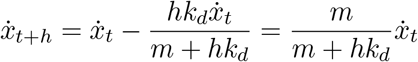

With *k_d_* now in the denominator, even an infinite damping gain is stable, corresponding to the damping force that completely eliminates the current velocity in a single timestep. This stability allows us to make PD controllers with extremely stiff gains.

**Fig 13.**
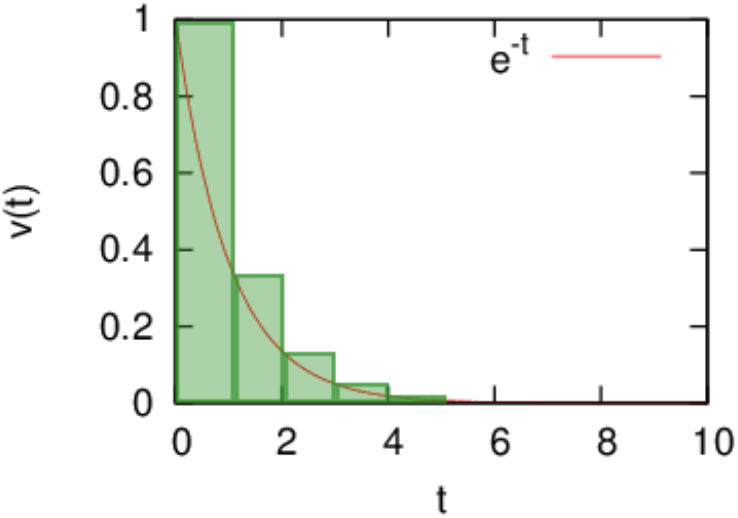
Explicit integration of damping forces is similar to the forward-method for approximating the area under a curve as a sum of rectangles. In this case it severely overestimates, leading to instability.

Stability is a nice property for a controller or simulator to have. We now show that the *α* and *β* terms added to the constraint equation change them into implicit springs. To see the correspondence between Eq. 10 and Eq. 14, we consider a constraint that keeps a point mass at the origin along a single dimension: *ϕ*(*x_t_*) = *x_t_*. The displacement function for this system has a trivial Jacobian: *J* = 1, meaning that *λ* = *f_c_*. Assuming that external forces are zero, *f_g_* = *f_u_* = 0, Eq. 10 simplifies to

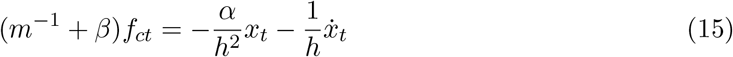

Assigning the *α* and *β* parameters^6^ to be, 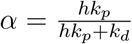, and 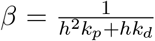, and isolating *f_ct_*, Eq. 15 reduces to the implicit spring equation Eq: 14:

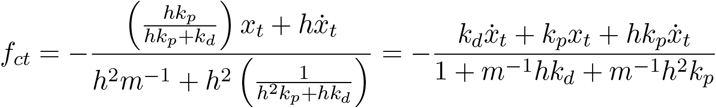

The consequence of this relationship is that every constraint in ODE can be thought of as an implicit spring. An important feature of this formulation is that the equations are solved simultaneously. When the implicit springs are solved simultaneously in the physics framework, the forces account for each other; without this change the system would be very fragile. Softening the constraints to springs makes it so that we can solve a system that would otherwise be over constrained. We can add more constraints than there are degrees of freedom.

#### Solving with Complementarity Conditions

For simplicity, we compress Eq. 10 down to ***Aλ*** = ***w***. When ***A*** is non-singular, we can solve for ***λ*** by inverting, or preferentially, using a fast, numerically-stable solver such as a Cholesky decomposition. Some constraints, however, come with additional conditions that need to be solved with extra machinery. In simulation literature, these are known as inequality constraints. For example, a contact constraint keeps two bodies from moving towards each other by defining an error function that is the separation of the contacting surfaces in the direction of one of the surface normals. If the surfaces are overlapping, then the error function has a negative value and a positive constraint force will accelerate the surfaces apart. This acceleration is as it should be. However, the linear system also applies forces to correct positive error; so the same constraint would also prevent the surfaces from separating.

The solution to this problem is to limit the amount of force available for satisfying the constraint. A contact constraint, in particular, limits the force to be non-negative. Contact friction constraints are limited on both sides to be proportional to the contact normal force. This limitation places upper and lower bounds on the constraint force variable: ***λ***_lo_ ≤ ***λ*** ≤ ***λ***_hi_, allowing constrained bodies to accelerate without bounds if the force necessary to hit the acceleration target falls outside of the limits. In ODE, the result is three possible conditions to satisfy a constraint:

1. ***a**_i_λ* = *w_i_* with *λ_i_* ∈ [*λ*_*i*lo,_ *λ*_*i*hi_],
2. ***a**_i_λ* > *w_i_* with *λ_i_* = *λ*_*i*lo,_ or
3. ***a**_i_λ* < *w_i_* with *λ_i_* = *λ*_*i*hi,_

where −∞ ≤ *λ_i_*_lo_ ≤ 0 ≤ *λ_i_*_hi_ ≤ ∞.

A linear solver cannot handle these extra conditions on the constraint forces. To solve this type of system, physics engines employ a mixed Linear Complementarity Problem (mLCP) solver. ODE offers two different solving methods for satisfying constraints under limited-force conditions. One method, known as Projected-Gauss-Seidel, solves constraints iteratively and accumulates the effects [56]. Iterative methods tend to be faster, but also tend to be inaccurate when the system is near-singular or ill-conditioned. Simulated humanoid systems, particularly with two feet on the ground, tend to behave badly with this faster solver. The slower, pivot-based method, follows the algorithm presented by Baraff [57]. Baraff’s method is still easily fast enough for our purposes.

Each row in matrix ***A*** represents a constraint. The corresponding values of ***w*** and ***λ*** represent a “target” acceleration along the degree of freedom constrained by that row and the generalized force used to achieve it. For the *i*^th^ row of ***A***, the diagonal element, *a_ii_*, behaves like the inverse mass of the constraint. A force, *λ_i_*, imposes an acceleration of *a_ii_λ_i_* = *w_i_* within the constraint error-space. The rest of the elements in a row of ***A*** encode the force’s effects on other constraint dimensions. A change in the *i^th^* constraint force *λ_i_* affects the *j*^th^ constraint space by accelerating it according to *δw_j_* = *a_ij_δλ_i_*. The order of the constraints is arbitrary and they can be rearranged as long as every row-swap is accompanied by the corresponding column-swap that maintains the proper symmetry.

Baraff’s solving algorithm (based on Dantzig’s simplex method) takes advantage of this arbitrary ordering by dividing constraints into different sets: a satisfied set *S*, a limited set *N*, and an unaddressed set *U*. All constraints fit into one of these categories. The first step in finding a solution is to reorder and satisfy all the unlimited constraints, without considering the rest, using a basic linear solver. The resulting system looks like

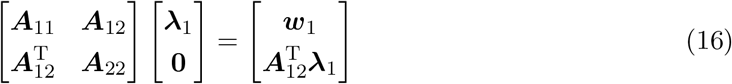

Set *S* holds the rows of ***A***_1*i*_. Set *U* holds the rest. At this point it helps to look at some figures to see what is going on. Each constraint’s target conditions can be represented as a piecewise line through force-acceleration space (Fig 14). We will call this multi-segmented line the target manifold for each constraint. Viewing constraints this way is another contribution of this work. The diagonal element of ***A*** associated with the constraint gives the slope of a line through the origin that represents the relationship between force (*λ*) and actual acceleration (*A_ii_* is the effective inverse-mass of the *i*^th^ constraint). The solver seeks to find a joint solution so that, for all rows of ***A***, the pairs of (*λ_i_, w_i_*) fall on the acceptable manifold. Forces from other constraints move the entire manifold up or down relative to the origin.

**Fig 14.**
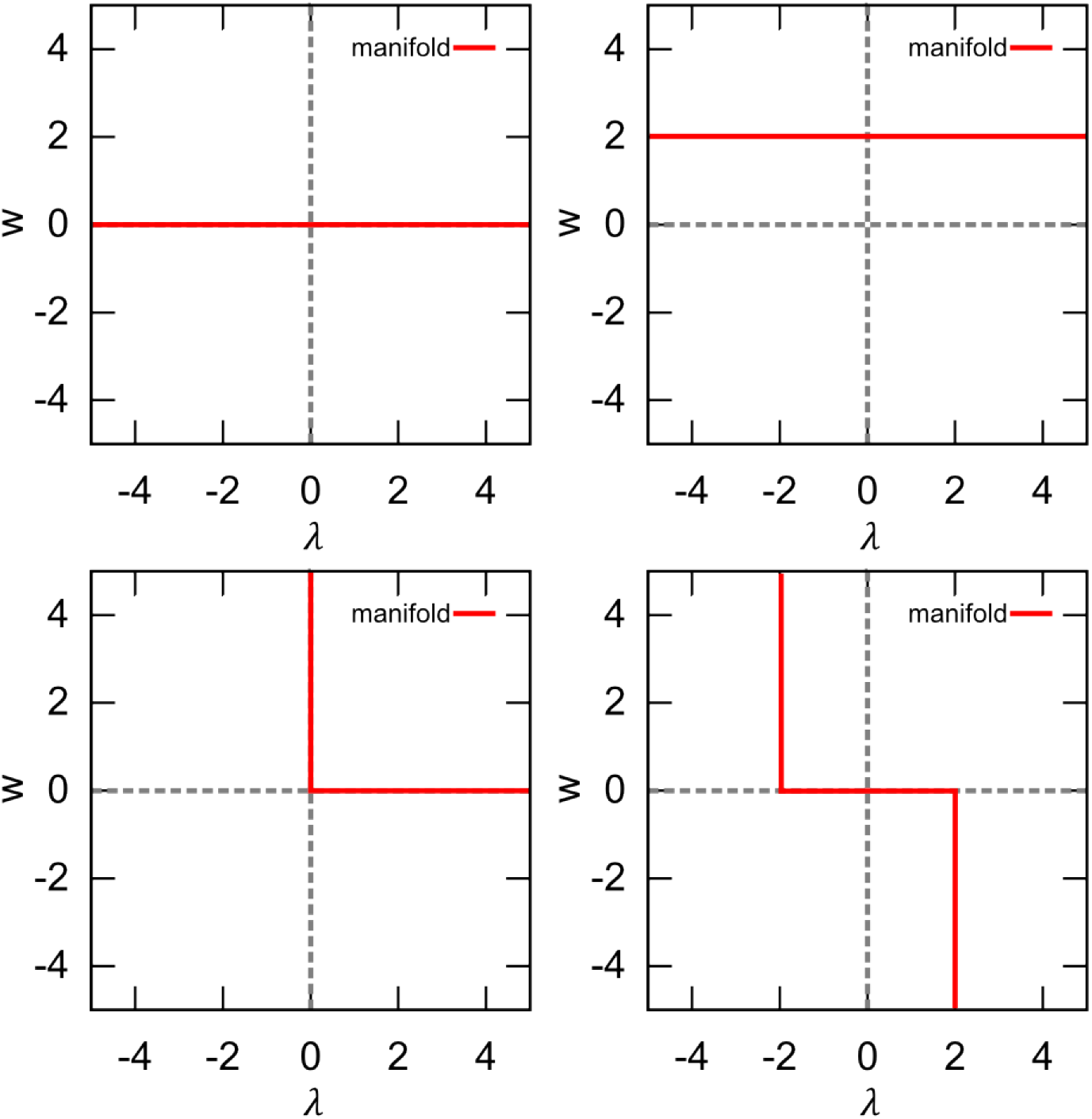
Each constraint on a single degree of freedom can be thought of as a monotonically decreasing, piece-wise linear target manifold through acceleration-force space.

The *β* parameter takes the horizontal portion of the target manifold and tilts it so that when bigger forces are used, there is a lower target acceleration. Hence the constraint is spring-like. The vertical portions of the constraint represent places where the constraint has hit its force limits. No additional force can be applied by that constraint; so the acceleration must be allowed to increase freely. Otherwise, the constraint would be “obligated” to apply more force to try to get closer to its target acceleration.

Constraints are addressed one-at-a-time. When dealing with ground contact force without softened constraints, once the solver found a sufficient force to keep a body from penetrating the ground, any remaining ground contact constraints would have nothing to do, resulting in inappropriate distribution of ground forces. With spring-like constraints, if one contact constraint supporting a body reaches its target force/acceleration, a second, redundant contact constraint will see whatever distance remains between the current acceleration and the target. Forces applied by the second constraint attempting to reach its target push the target manifold of the first constraint toward the origin. The force required to achieve the first constraint’s target decreases until the forces balance appropriately. The balancing forces make it possible to more accurately compute inverse dynamics forces.

The algorithm for solving the mLCP progresses through each unaddressed constraint, one at a time, and finds the change in forces that will satisfy the new constraint without moving any of the current constraints off their piecewise target. Each iteration of the algorithm draws a new constraint from the unaddressed set *U* and addresses the change in force, *λ*, that will satisfy the new row without pushing any previously addressed rows off their manifold, until the new row can be added to *S* or *N*. In the process other rows may change between sets *S* and *N*, but each row remains on its target manifold in acceleration/force space.

Consider this partitioned matrix:

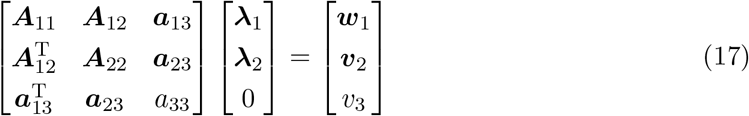

Adding a new force, *λ*_3_, will change the accelerations of the other constraints. Accelerations of constraints at their limit are allowed to change, but those in set *S* must remain at their target. So we find the *δλ*_3_ that moves *v*_3_ toward *w*_3_ and find the simultaneous *δ**λ***_1_ that keeps constraints in *S* satisfied. The constraint force takes the largest step that will not push any row out of its set. This step will either satisfy the constraint or move another constraint to an intersection point on its manifold. We then pivot the sets around and continue until all of our rows are in *S* or *N*. For additional detail, see [57].

Recognizing that the solver deals with each constraint target as a piecewise linear manifold provides useful insight into how the simulation mechanism can be improved. One obvious extension is to increase the number of linear segments in the target manifold beyond three (Fig. 15). This innovation becomes obvious when constraints are considered as target manifolds rather than Lagrange multipliers. With a multi-segment target manifold, it is possible to create a spring-like constraint that is loose near its setpoint, but then becomes stiffer.

**Fig 15.**
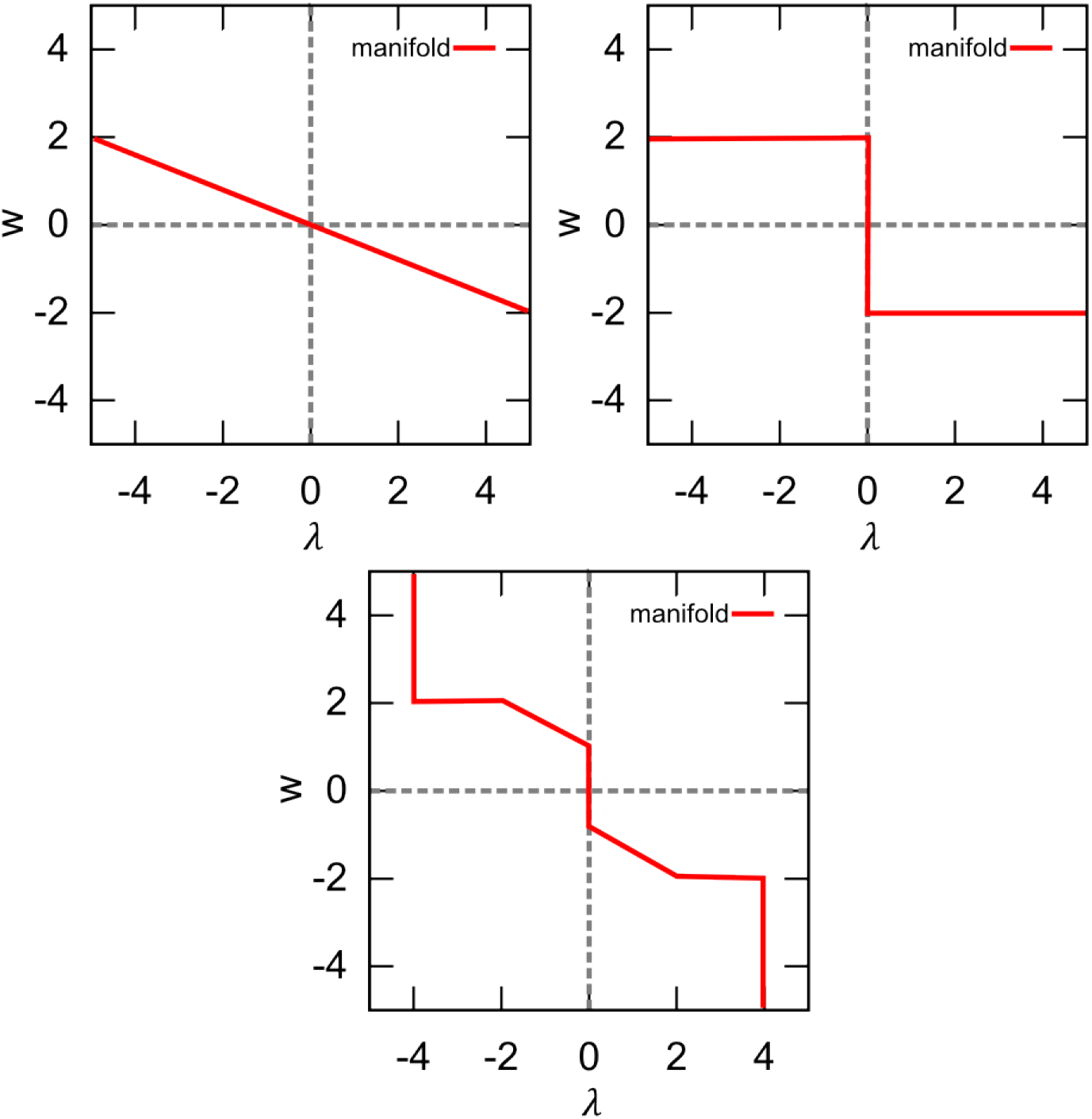
Adding a small value to the diagonal elements of the projected inverse mass matrix turns the constraint into a spring. Viewing constraints as piecewise linear targets provides insights into how to make more complicated constraints consisting of additional piecewise segments.

We can make spring constraints that get more or less stiff as additional force is required. We can also introduce constraints with “deadzones” in their PD control (Fig. 15). This type of constraint is particularly interesting because it allows us to introduce controllers that only come into play when a dimension of interest drifts out of an acceptable range. This type of controller takes inspiration from the idea of “uncontrolled manifolds” in human motor control theory [58]. With this constraint acting as a controller, if a perturbation will not hurt performance, the controller does nothing.

From deadzone controllers, we can introduce novel constraints with secondary targets. A constraint whose forces and accelerations fall within acceptable tolerances has flexibility to “help” another constraint that has reached its limit. For example, we can specify a target range for the knee, hip, and ankle joints of a simulated character. When these leg joints fall within their stated ranges, they can be allowed to pursue a secondary goal such as keeping the torso upright or at a given height. This type of constraint can serve as a method for reducing the need for unrealistic residual forces. Removing residual forces implies deviating from original kinematic data. Constraints with secondary targets make it intuitive and clear how this deviation will occur can be extremely beneficial when using the simulation engine for analyzing or synthesizing movement data. We have created and submitted code for allowing controller constraints with a deadzone in acceleration space. ^7^

### S2 Appendix

The model consists of *n_b_* rigid bodies connected by *n_j_* joints. In this case, each joint consists of three to five constraints. Each joint connects two rigid bodies with anchor points (center of rotation) defined in the reference frame of both bodies. The joint constraints keep the anchor points relative to the two bodies together in the global frame. If bodies *b_j_* and *b_k_* are connected, a joint constrains them together at a common point. The joint anchor relative to body *b_j_* is 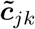. The anchor for body *b_k_* is 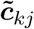. The joint constraint drives these points together in the global frame, creating three constraint rows:

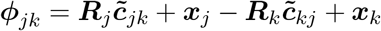

The locations of these anchor points determine the segment dimensions (bone lengths) of the character model.

Markers, each assigned to a specific rigid segment, represent a point on the human body. We seek anchor points that allow markers to remain approximately stationary relative to their assigned body segment. It is generally impossible to precisely find such a configuration (without creating an unreasonable number of body segments) because of soft-tissue artifacts (STAs). Skin and joints are not rigid. They stretch and give as muscles pull the bones. Modeling the body in maximal coordinates provides a way to model STAs explicitly.

Given a pre-defined model topology and markers assigned to specific model segments, we seek to find the joint anchor points between segments and the marker attachment points relative to the model segments. If the *i*^th^ marker is assigned to the *j*^th^ rigid body (***p**_i_* → *b_j_*) at relative point 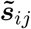, we model the marker’s attachment as a three dof constraint:

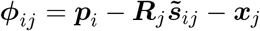

The process models markers from an arbitrary point in time as infinite point masses. As bodies of infinite mass, constraint forces do not affect the markers’ trajectories but only the bodies they are anchored to. Initially, markers are anchored at 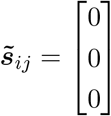. This mapping attaches the marker to body *b_j_*’s center of mass.

This mapping is a very rough estimate of the marker attachment points on the model segments, but it is sufficient because of the flexible nature of constraints in the simulation software. Setting the *CFM* parameter of the marker constraints to *β* = 10^*−*3^ and setting the model joint constraint *CFM* to *β* = 10^*−*5^ makes the body segments hold together tightly, while still allowing the markers to pull the body into shape. Several timesteps of simulation allow the model to relax to a fixed pose. We then take the markers in their current configuration and reattach them to their respective segments. Relaxing the marker attachments this way improves the fit for this particular frame of marker data. Iteratively repeating this process with multiple frames of marker data, we therafter update the marker attachment points by some learning rate, 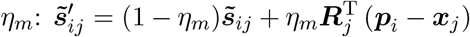. Gradually updating attachment points, using different frames of data, effectively descends the error gradient of the marker positions relative to the body:

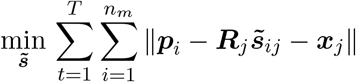

The decrease in marker error affected by model dimension error. Conveniently, joint anchor constraints behave the same as the marker attachment constraints. With an arbitrary frame of marker data and using a marker *CFM* of *β* = 10^*−*4^, if the markers constraints cannot be satisfied, they will pull the joint anchors apart slightly. For each joint we find a new common anchor point in the global frame by taking the average between the two unsatisfied anchor points that the joint constraint is trying to pull together. We then move the anchor points toward that point according to learning rate *η_l_*:

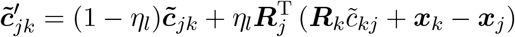

For any frame, errors will cause the markers to stretch from their attachment points and joint anchor points to stretch apart from each other. Both marker attachment points and the joint anchors can be updated simultaneously to decrease the error for that frame. However, the local solution that perfectly satisfies one frame may make another frame worse. This step presents an evident gradient descent approach to finding the joint anchors and marker attachments: using several frames, compute an average adjustment to the marker attachments, and joint anchors that reduce the error. Make the adjustment to both anchors and attachments and then iterate. It may be advisable to employ the standard machine learning practice of a validation set to ensure that the error continues to decrease and avoid overfitting. This technique relies on spring-like constraints made possible in maximal coordinates.

Although this method could easily be automated, in practice, the research did not rely on very many different models and so the system uses a mechanism for relaxing the marker attachment points and joint anchors with the click of a button in the graphical user interface (Fig. 1). With a new data set, a handful of iterations proved sufficient to produce a reasonable model with marker attachments that fit the data well enough for further analysis. This algorithm does not address joint limits on a range of motion. These can also be learned [44], but in our case, the range of motion for each joint is set *a priori*. After determining segment lengths, we set other segment dimensions as appropriate to fit against the markers. Mass properties for each segment assume uniform density by volume.

## Acknowledgments

Thanks to John Matthis and Leif Johnson for helpful discussions. This research was supported by NSF grant CNS1446578

## Footnotes

1 OpenDE: http://www.ode.org/

2 MuJoCo http://www.mujoco.org/

3 The HDM mode: https://github.com/EmbodiedCognition/QtVR

4 HDM UI Demo https://youtu.be/ASs4Wo5PQcM

5 CMU Graphics Lab Motion Capture Database: http://mocap.cs.cmu.edu/

6 These values are presented without derivation in the ODE user-manual: http://ode-wiki.org/wiki/index.php?title=Manual:_All#How_To_Use_ERP_and_CFM. Note that our formulation of *β* has an extra *h* in the denominator which is added automatically by ODE.

7 Full implementation of secondary targets for constraints is still in progress. It promises to be useful for creating intelligent constraint-based controllers.

